# Frontal cortex norepinephrine, serotonin, and dopamine dynamics in an innate fear-reward behavioral model

**DOI:** 10.1101/2023.11.27.568929

**Authors:** Jen-Hau Yang, Emily L. Burke, Aakash Basu, Rong-Jian Liu, Stephanie M. Staszko, Abigail L. Yu, Jocelyne Rondeau, Samira Glaeser-Khan, Yizhou Zhuo, Jiesi Feng, Yulong Li, Alicia Che, Alfred P. Kaye

**Author notes:** Corresponding author: Alfred P. Kaye. Equal Contributions (Jen-Hau Yang, Emily L. Burke).

## Abstract

Animals must survive by foraging for food in an uncertain and dangerous world. Neural circuits for naturalistic decision-making under these conditions must balance competing motivational demands for approaching reward and avoiding danger. To enable flexible switching between motivational states, neuromodulators are released and alter neural excitability and plasticity. The question of how neuromodulators encode motivational state is thus fundamental to systems neuroscience, yet the dynamics of neuromodulators during naturalistic decision making are not fully understood. Here, we developed a naturalistic approach/avoidance task in mice involving a tradeoff between seeking reward versus safety in the presence of looming predation risk. We utilized fiber photometry, computational behavior tracking, and local pharmacology in this task. Silencing medial prefrontal cortex (mPFC) reduced looming defensive behaviors. Moreover, by using fiber photometry combined with GPCR-based sensors, we found that cortical norepinephrine (NE) plays a more prominent role in encoding looming threats while dopamine (DA) represents reward and threat. In contrast, serotonin (5HT) dynamic negatively correlates to both emotional valences. To begin to understand neuromodulatory interactions, we used *ex vivo* slice physiology to understand 5HT impact on spontaneous firing of locus coeruleus NE neurons. In conclusion, monoamines such as NE, DA, 5HT can converge in their encoding of naturalistic motivated behaviors as well as dissociate from one another. By utilizing this novel innate reward-threat task, we can better understand neurochemical signaling events during natural behavior, and may contribute to the understanding of neural mechanisms underlying emotional decision-making and its implications for psychiatric disorders.

## Introduction

Anxiety and posttraumatic stress disorder (PTSD) are characterized by a bias towards the excessive appraisal and reactivity to threat. A significant challenge in the translational modeling of psychiatric disorders is to connect moment-to-moment decision-making about risk to the states of prolonged maladaptive anxiety. One promising direction has been to study situations in which subjects must approach reward while avoiding harm (Ito & Lee, 2016; Kirlic et al., 2017; La-Vu et al., 2020; Chen et al., 2022). The medial prefrontal cortex (mPFC) integrates sensory evidence about the world with internal motivational state during approach-avoidance decision-making. In humans, neuroimaging studies of risky decision-making and genomic studies of PTSD have both implicated mPFC as a critical node.

A limitation of our understanding of how subjects navigate approach-avoidance motivational conflict is that many studies have used learned reward and fear signals (such as tone-shock pairing) rather than naturalistic danger signals. Naturalistic approach-avoidance behavioral paradigms might enable more ethological mappings from circuits to behavior (Silva et al., 2016; Evans et al., 2019). To study the neural mechanisms underlying this process, aversive stimuli such as foot shocks, predator odors, startle tone, and looming have been widely used in rodent research (Adolphs, 2013; Ganella & Kim 2014; Bedoya-Perez et al., 2019). The looming predator stimulus mimics visual cues indicative of aerial predators to reliably induce rapid innate fear responses including freezing and escaping (Yilmaz & Meister, 2013; Evans et al., 2018). Looming recruits distributed subcortical and cortical circuits, including amygdala, thalamus, superior colliculus, and mPFC (Zhao et al., 2014; Wei et al., 2015; Shang et al., 2015, 2018; Hu et al., 2017; Evans et al., 2018; Salay et al, 2018; Vander Weele et al., 2018; Fratzl et al., 2021).

Neuromodulators also modulate innate fear responses (Huang et al., 2017; Li et al., 2018; Zhou et al., 2019; Barbano et al., 2020), raising the question of how their moment-to-moment dynamics may affect naturalistic approach-avoidance behavior. Neuromodulators such as norepinephrine (NE), dopamine (DA), and serotonin (5HT) have distinct and region-specific effects on switching between motivational drives and project to mPFC. For example, NE represents exploration-exploitation tradeoffs or uncertainty within threat (Yu & Dayan, 2005; Sciolino et al., 2022; Basu et al., 2024), DA canonically represents reward prediction errors and motivated behavior (Schultz, 2010, 2016; Salamone & Correa, 2012; Zald & Treadway, 2017; Salamone et al., 2018), and 5HT functionally modulates emotional motives (Marcinkiewcz et al., 2016; Huang et al., 2017; Ren et al., 2018). Among subcortical and cortical transmission networks, the prefrontal cortex stands out as a critical region that receives convergent neuromodulatory inputs to process complex cognitive functions (Cools & Arnsten, 2022). mPFC has been shown to encode sensory salience by incorporating neuromodulatory inputs (Ventura et al., 2008; Kim et al., 2018; Ritter et al., bioRxiv). However, it remains unknown how rapid neuromodulator dynamics in mPFC are organized during naturalistic motivational conflict and how they distinctly they represent emotional valence, arousal and behavioral switching.

In this study, we developed a naturalistic approach-avoidance conflict task involving looming predator stimulus and food reward and characterized adaptations in behavior over days. We used local pharmacology to demonstrate a role for mPFC in this behavior, then assessed the dynamics of neuromodulators in freely moving mice in the task using fiber photometry with GPCR activation-based (GRAB) sensors. The GRAB sensors are genetically encoded neuromodulator indicators, enabling direct measurement of evoked neuromodulator release *in vivo* (Sun et al., 2018, 2020; Feng et al., 2019; Wan et al., 2021; Zhuo et al., bioRxiv). Comparative analysis of DA, NE, and 5HT across cohorts showed distinct encoding of innate reward and threat during aspects of the task, and slice electrophysiology and pharmacological manipulations further characterized 5HT-NE interactions relevant to emotional conflict.

## Results

### Approach/avoidance behavior in an innate emotional conflict task

To investigate the motivational conflict between foraging for reward and environmental threats, we developed a naturalistic approach/avoidance task in mice (**Figure 1A**). In this task, freely moving mice were allowed to explore in a rectangle arena for reward readily upon licking the spout (i.e., approach, **Figure 1B**) or to flee and hide in a red-tinted shelter from visual looming stimuli (i.e., avoidance, **Figure 1C**). The looming stimulus was displayed as a video of a quickly expanding black disk above the arena to simulate approaching aerial predator, which reliably caused innate fear responses (Yilmaz & Meister, 2013; Evans et al., 2018). Looming was randomly presented twice in each of 5 daily sessions. All behaviors were recorded throughout each 20-25 min session and analyzed offline with Matlab and Deeplabcut (**Figure S3A&B**, Mathis et al, 2018). At the first looming stimulus, 96% of mice immediately exhibited classic fear responses (freezing or escaping), but this percentage decreased to 52% at the 10th looming stimulus (**Figure 1D; Figure S1**). In contrast, the number of licks within the first 180 seconds increased significantly across days (F(4, 120) = 4.084, *p* < 0.005), and multiple comparisons with adjusted *p* value further indicated significant differences at Day 1 versus Day 3 and Day 1 versus Day 4 (both *p* < 0.05, **Figure 1E**). There was no sex difference in total licks between male (178.7 ± 38.6) and female (225.1 ± 51.1, p = 0.17, **Figure S3A**). In this conflict task, mice showed adaptive behavior in a manner of decreased looming-induced fear responses and increased number of licks, indicating a natural flexible switch of behavior from more escaping to more reward seeking (**Figure 1F**). To determine how looming threats directly impact licking behavior, we analyzed the events where the looming was displayed while the mouse was licking or not (**Figure 1G**). The area under the curve (AUC) of each daily psychometric function in likelihood toward the next lick increased across days, suggesting an elevated tendency to overcome the looming threat over time. These findings collectively demonstrated an adaptive behavior in mice performing the conflict task, akin to the vigorous flexibility seen in animals foraging under predatory threats in the wild.

**Figure 1.**
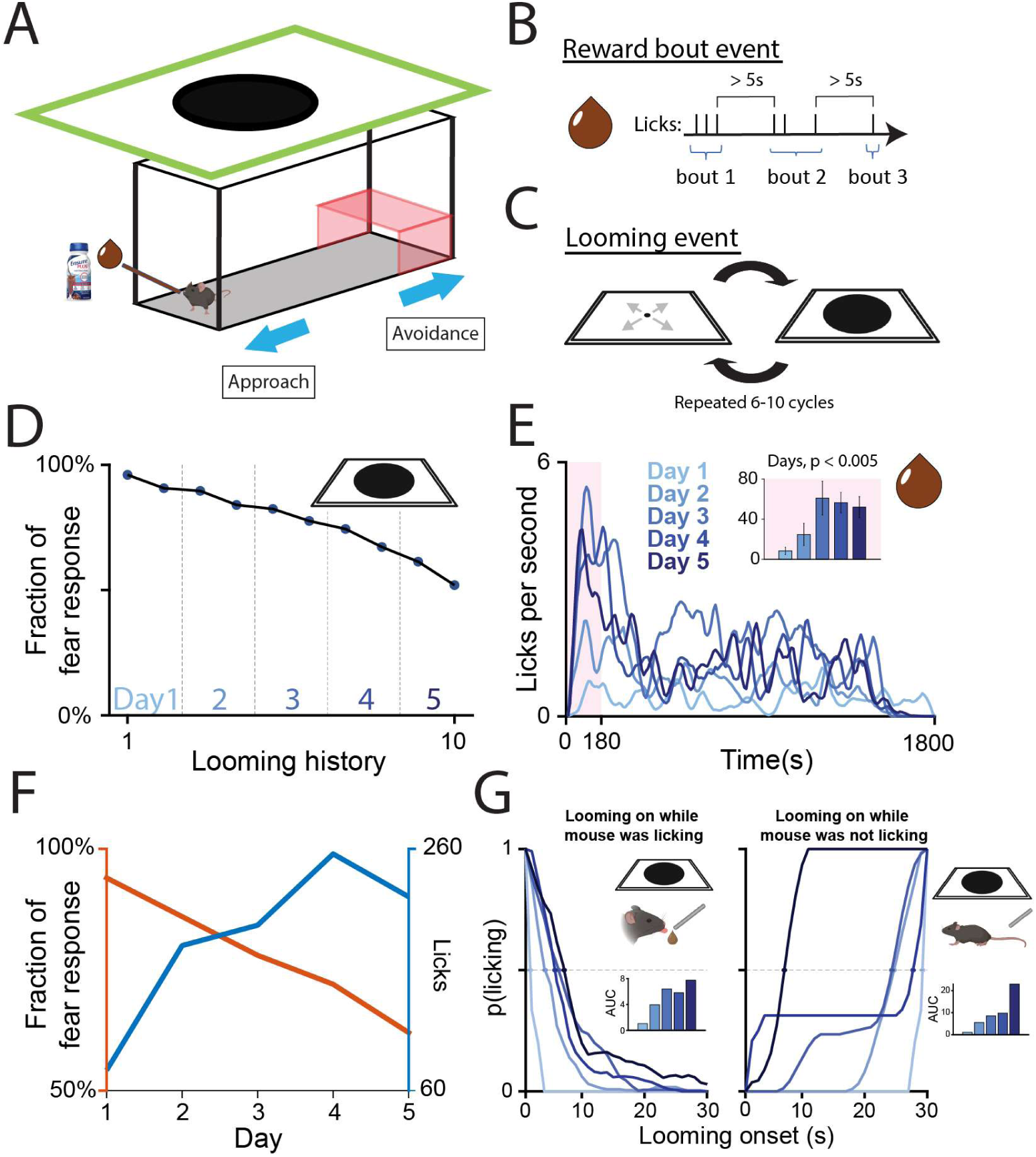
A naturalistic motivational conflict task which assesses reward foraging and innate fear responses. **A**: Diagram of the approach/avoidance conflict task setting. Each mouse is allowed to freely lick for reward (Ensure), but with unexpected looming stimuli present from above. **B**: Event of a reward bout, which is defined as a train of continuous licks within 5 seconds from last lick. **C**: Event of the Looming stimulus, which is displayed as a quick expanding black disk mimicking an approaching aerial predator. **D**: Percentage of fear responses induced by looming stimuli across five daily days (n = 25). **E**: Lick distribution (in Hz) across 5 daily sessions. Inset figure indicates a one-way ANOVA analysis with the average licks (±S.E.M.) within the first 180 seconds (pink background) each day. **F**: Adaptive behavior indicated by a multiple scale of fear response and number of licks. **G**: Event of looming stimulus was presented while the mouse was licking (left) versus not licking (right panel). Inset figures represent the area under the curve (AUC) each day.

### Prefrontal NE encodes looming threats, and DA encodes reward bouts

Prior studies demonstrated that both reward and aversive decision-making involves prelimbic mPFC (Capuzzo et al 2020; Fernandez-Leon et al 2021), so we hypothesized that innate reward and aversive decision-making would also involve mPFC. Midline thalamic projections to mPFC modulate fear-related responses to looming (Salay et al 2018). To verify mPFC’s activity is necessary for looming-induced fear responses, muscimol, a GABA agonist, was intracranially infused into the prelimbic region of the mPFC to inhibit neuronal activity. Following infusion, mice underwent the innate fear-reward task, and their behavior in response to the looming stimulus was quantified (**Figure 2A**). Behavior typically fell into one of three categories: freezing, escaping to the shelter, or no response. Muscimol infusion caused a decrease in fear-related behaviors (i.e. freezing and escaping), and in turn increased no response behavior in response to the looming stimuli (**Figure 2B**, GLMM: OR = 0.15, 95% CI [0.03, 0.88], p = 0.037). Consistent with these findings, we also observed that exposure to looming induced cFos in prelimbic mPFC (**Figure S2**).

Multiple neuromodulators influence looming behavior; given mPFC’s role in opposing emotional valence behaviors we sought to directly measure *in vivo* NE and DA release in mPFC in mice over the course of repeated exposure to in our reward-looming task. In this study, cortical NE and DA dynamics were measured in mPFC by using G-protein-coupled receptor activation based (GRAB) sensors (GRAB_NE_: AAV9-hSyn-NE2h; GRAB_DA_: AAV9-hSyn-DA3m), which are fluorescent indicators of neuromodulator release (Sun et al., 2018, 2020; Feng et al., 2019, **Figure 2A&B**). Time-locked looming- and reward-related neurotransmitter fluorescence traces were compiled by aligning the event onset time (t = 0, **Figure 2**), and a one-way repeated-measures ANOVA analysis for event-related NE df/f revealed a significant main effect in looming events across days (**Figure 2D**, F(4, 100) = 2.939, *p* < 0.05, multiple comparisons indicated significant difference between Day 1 vs. Day 4, *p* < 0.05) and reward-bout events (**Figure 2G**, F(4, 92) = 5.211, *p* < 0.001, and significant Day 2 vs. Day 4, *p* < 0.05; and Day 2 vs. Day 5, *p* < 0.001). On the other hand, DA dynamics showed no day main effect in looming (**Figure 2J**, F(4, 46) = 0.2839, p = 0.89), but a significant main effect in reward bout (**Figure 2M**, F(4, 42) = 3.748, *p* < 0.05, and significant Day 1 vs. Day 5, *p* < 0.05). Overall, the averaged mPFC NE dynamic plays a more prominent role in encoding looming threats over reward bouts (**Figure 2E&H**), whereas mPFC DA encodes reward bouts, especially in early days (**Figure 2K&N**).

**Figure 2.**
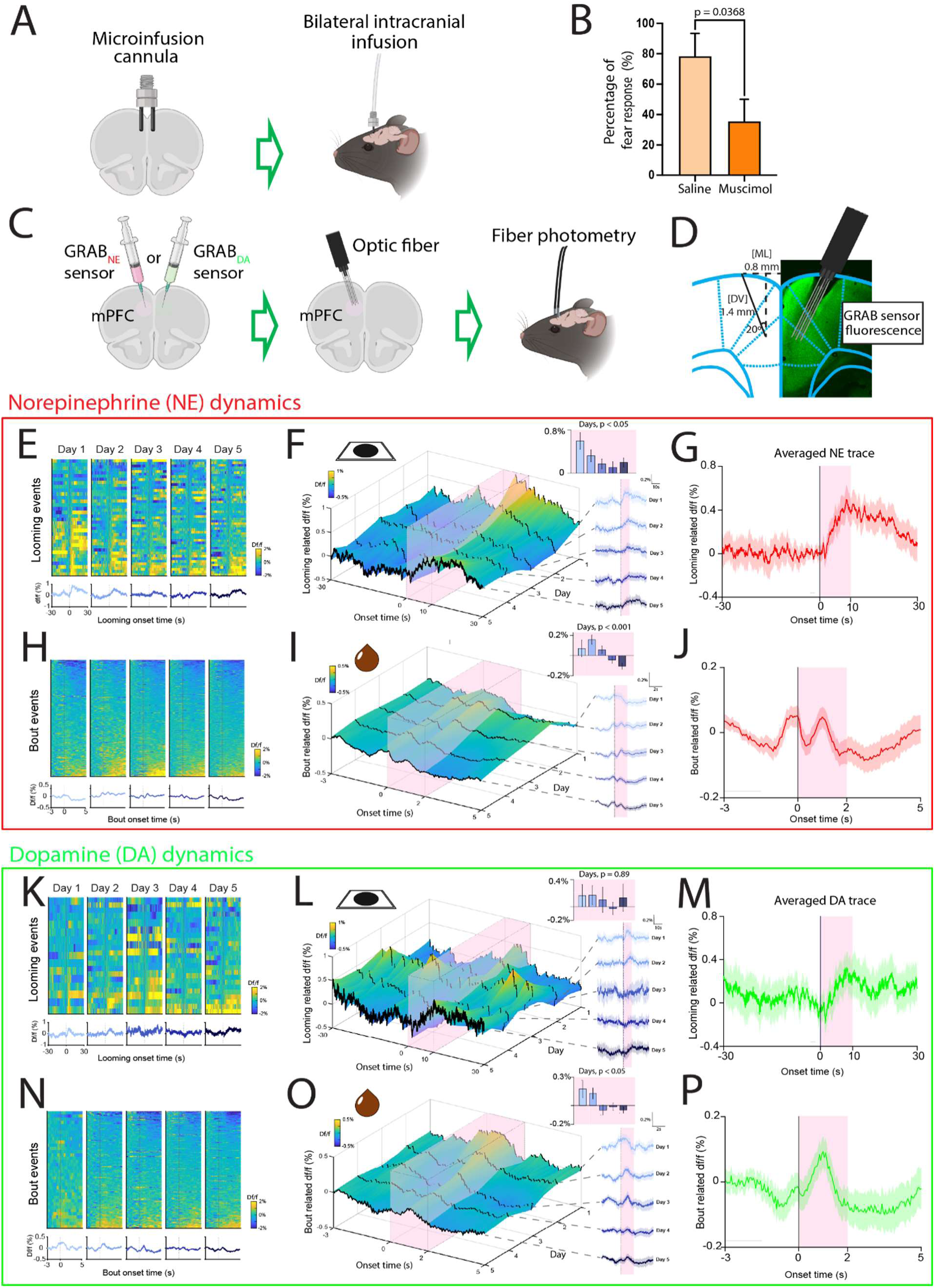
Local pharmacology and fiber photometry with GRAB sensors for measuring *in vivo* event-dependent neuromodulators dynamics. **A**: Diagram of microinfusion cannula placements targeting mouse mPFC. **B**: Percentage of looming trials in which mice exhibited a fear response (either escape or freezing) after local infusion of either saline or muscimol. **C**: Diagram of bilateral GRAB sensor injections and optic fiber implements targeted mouse mPFC. **D**: Example histology photo of GRAB sensor expression and fiber track with a 20° angle. **E-J**: Event-related norepinephrine (NE) dynamics to looming stimuli and reward bouts. **E**: Heatmaps (top) of looming-induced individual NE traces 30 seconds before and after event onset (t = 0) and average (bottom) across days. **F**: Three-dimension representation of day-by-day looming-related NE dynamic change across each daily session (right). Shaded area indicates the 95% of confidence intervals. Inset figure indicates a one-way ANOVA analysis of mean NE activity (±S.E.M.) within the 0 to 10s window (pink background). **G**: Averaged NE trace of all looming events. **H**: Heatmaps of reward-bout-induced individual NE traces (top) and average (bottom) across days. **I**: Three-dimension representation of bout-related averaged NE dynamic across each daily session (right). Inset figure indicates a one-way ANOVA analysis of mean NE activity within the 0 to 2s window. **J**: Averaged NE trace of all bout events. **K-P**: Event-related dopamine (DA) dynamics to looming stimuli and reward bouts.

### Prefrontal NE is activated during threat exposure

Looming-evoked freezing and escaping have been previously shown to be promoted by divergent pathways (Salay et al., 2018; Shang et al., 2018), it is thus inferred that the same threat signal may trigger dissociative neuromodulatory dynamics corresponding to different behavioral response types. In this task, looming-evoked fear responses were categorized as freezing, escaping, and no response (**Figure 3A&B**). Overall, mice predominantly showed more freezing (57%) than escaping (21%) and no response (22%), and increased no response was associated with shorter latency to next lick (**Figure 3C**). Also, we found that male mice tend to freeze more to the looming stimuli compared with female mice (**Figure 3D**). By synchronizing photometry and behavioral tracking data, we next assess neuromodulatory effects on the different response types (**Figure 3E&F, S2**). Within a 15s window after looming onset, a one-way repeated-measures ANOVA revealed a significant response-type main effect in mPFC NE activity (F(2, 207) = 4.901, *p* < 0.01) and in DA (F(2, 99) = 3.231, *p* < 0.05). In NE activity, multiple comparisons further indicated significant differences between freezing and escaping (*p* < 0.01). In contrast, multiple comparisons in DA activity revealed no significant differences in freezing versus escaping (p = 0.17) or freezing versus no response (p = 0.08). mPFC NE dynamics were more sustained in freezing mice, who also have more prolonged exposure to looming compared to mice that escaped to shelter (Figure 3E&F). There were subtle differences in NE activity over days, with slightly increased NE in day one during escape and freezing responses to the looming stimuli (Figure S3A). Differently, there were little to no differences in DA activity over days for any of the looming-induced behaviors (Figure S3B; effects over days and sex differences Figure S5-S6). In addition, when propranolol, a ß-adrenergic receptor antagonist, was locally infused into the mPFC, it specifically decreased freezing but not escape behaviors in response to the looming stimuli (**Figure S2F**, GLMM: OR = 0.16, 95% CI [0.03, 0.89], p = 0.0376)

**Figure 3.**
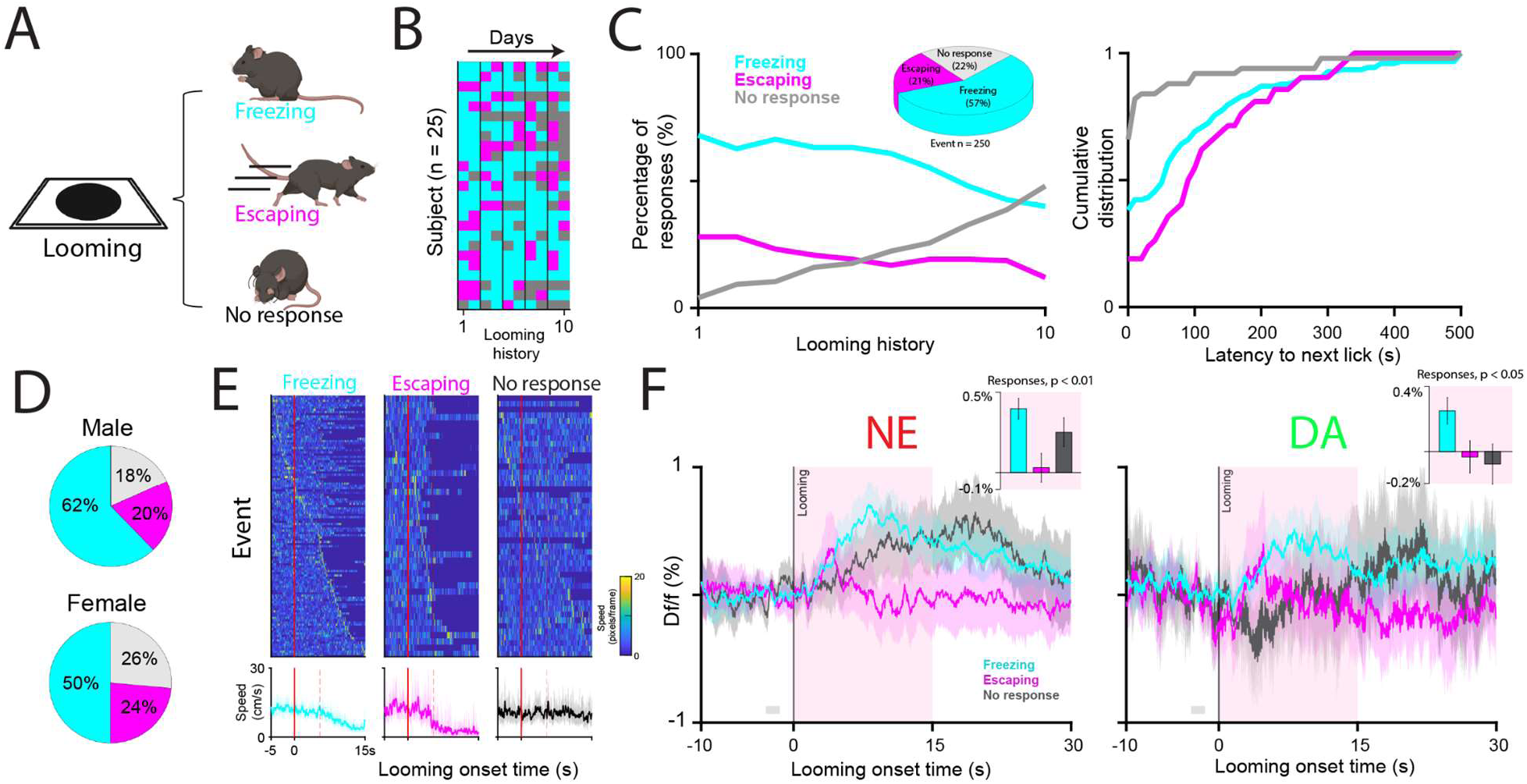
Dissociative neuromodulatory dynamics between looming-induced freezing and escaping behavior. **A**: Diagram of looming-induced responses, including freezing, escaping, and no response. **B**: Heatmap representation of individual mouse (n = 25) response across looming stimuli. **C**: (Left) Percentage of freezing, escaping, and no response over 10 looming stimuli. Piechart represents the overall ratio of behavioral responses. (Right) Cumulative distribution of latency to next lick after freezing, escaping, and no response behavioral subtypes. **D**: Percentage of looming-induced behavior responses between male (n = 15) and female (n = 10) mice. **E**: Heatmaps of looming-induced freezing (left), escaping (middle), and no response (right) on speed changes in a −5 to 15s window from looming onset (red line). (Bottom) Averaged speed (±95% confidence interval) from each day. **F**: Norepinephrine (left) and dopamine (right) dynamic changes associated with looming-induced responses. Shaded area indicates the 95% of confidence intervals. Inset figure represents a one-way ANOVA analysis of averaged df/f (±S.E.M.) within the 0 to 15s window (pink background).

### Prefrontal 5HT activity is negatively correlated to looming threats and reward

Next, we sought to investigate the role of mPFC 5HT in the conflict task. We hypothesized that looming threats may reduce mPFC 5HT activity, based on prior findings that looming decreased activity in DRN 5HT neurons (Huang et al 2017). In order to enhance the motivational conflict, we added a pure-tone cue which indicates reward availability and potential looming stimuli, creating more confined conflict epochs to the same approach/avoidance task paradigm (**Figure 4A**). This ensured that reward availability was solely during potential threat presentation instead of free availability as in previous experiments. In this 8-day task, mice were trained to associate the tone-on to reward in the first 4 days (training day T1 to T4), and the looming were added through day 5 to 8 (looming day L1 to L4, **Figure 4B&C**). Behaviorally, compared to well-trained day T4, mice licked significantly fewer on L1 (*p* < 0.05, **Figure S7**) while keeping their high hit rate unchanged across all four looming days (all *p* > 0.05), suggesting that mice were able to maintain the accuracy in obtaining reward under looming threats (**Figure 4D**). One-way repeated-measures ANOVA revealed no significant day-by-day main effects on 5HT dynamic changes during looming (Figure 4F, F(3, 38) = 1.260, p = 0.30) or bout events (Figure 4I, F(7, 75) = 1.331, p = 0.25), though 5HT showed consistent time-locked decreases to both looming and reward on individual trials that did not change over time (Figure 4G&J). Regarding looming-induced behavioral responses, a linear mixed-effects model with mouse as a random intercept revealed a significant response-type effect on 5HT release (χ^2^(2) = 10.96, p = 0.004; 84 trials, 11 mice), with pairwise comparisons showing a significant difference between freezing and escaping (b = 0.0018, p < 0.001, Figure 4K). As with NE and DA, there were exploratory differences in 5HT responses to looming across days when examined by behavior type (Figure S8-S9). Lastly, we sought to investigate how looming stimuli and licking behavior affect the bout signal in 5HT (**Figure 4L**). Before looming stimuli were introduced, the pure reward signal encoded by mPFC 5HT showed an immediate decrease after reward bout onset and rebounded to baseline. Interestingly, after looming stimuli were added, the bout-related signals showed divergent patterns according to whether the mouse kept licking or not (**Figure 4L**). Together, these results suggested that mPFC 5HT is negatively correlated to both negative and positive valences in a behavior- and time-dependent manner.

**Figure 4.**
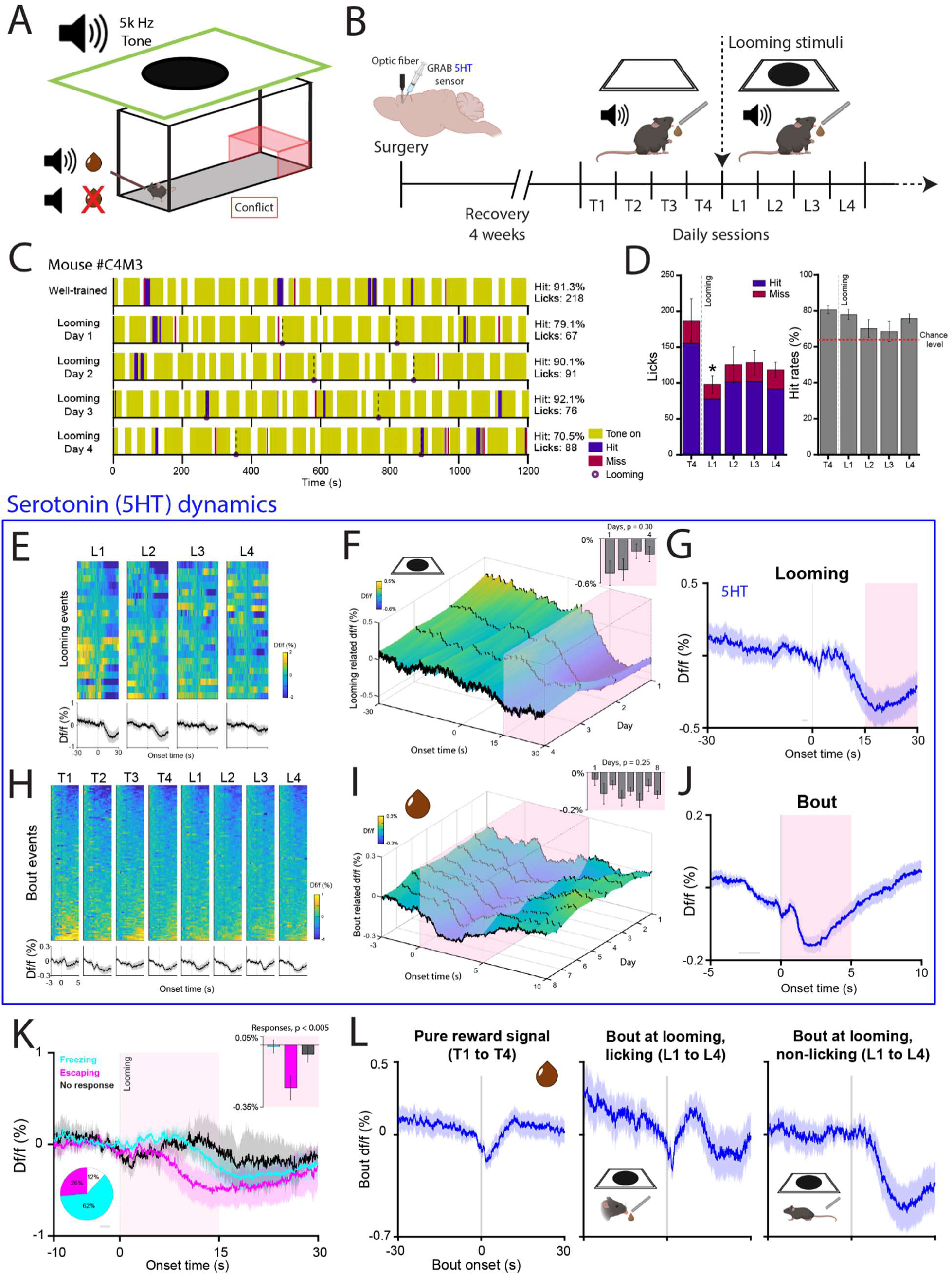
Serotonin (5HT) dynamics in a tone version of conflict task. **A**: Diagram of a tone component added in the approach/avoidance conflict task. Tone-on indicates reward availability and potential looming stimuli. **B**: Experimental schema of the task. After surgery and recovery, the behavioral task is divided into 4 training (T1 to T4) and 4 looming days (L1 to L4). Looming stimuli are introduced only on L1 to L4. **C**: Example responses of a mouse performing the task in well-trained (T4) and looming days (L1 to L4). Licks during tone-on are recorded as ‘hits’, otherwise ‘misses’ during tone-off period. **D**: Average licks (left) and hit rates (right) on T1 and L1 to L4. *: *p* < 0.05 planned comparisons to T4. **E-J**: Event-related serotonin (5HT) dynamics to looming stimuli and reward bouts. **E**: Heatmaps (top) of looming-induced individual 5HT traces 30 seconds before and after event onset (t = 0) and average (bottom) across 4 looming days. Shaded area indicates the 95% of confidence intervals. **F**: Three-dimension representation of looming-related 5HT dynamics over days. Inset figure indicates a one-way ANOVA analysis of mean 5HT activity (±S.E.M.) within the 15 to 30s window (pink background). **G**: Averaged 5HT trace of all looming events. **H**: Heatmaps of reward-bout-induced individual 5HT traces (top) and average (bottom) across days. **I**: Three-dimension representation of bout-related averaged NE dynamic across each daily session. Inset figure indicates a one-way ANOVA analysis of mean 5HT activity within the 0 to 5s window. **J**: Averaged 5HT trace of all bout events. **K**: 5HT dynamics associated with looming-induced freezing, escaping, and no response. Inset figure represents a one-way ANOVA analysis of averaged df/f (±S.E.M.) within the 0 to 15s window (pink background). **L**: Averaged 5HT traces (±95% confidence interval) of pure reward signal (T1 to T4, left). 5HT activity to bout events between mouse keep licking when looming was on (middle) and mouse quit licking (right).

### NE, DA, and 5HT showed distinct encodings of negative and positive valences

To better understand the neuromodulatory effects of NE, DA, and 5HT in totality, we compared their time-locked dynamics on looming and reward bout events across groups (**Figure 5**). In looming-evoked df/f, a one-way repeated-measures ANOVA revealed significant neuromodulator main effect (F(2, 44) = 58.93, *p* < 0.0001) on immediate activity (t = 0.5 to 1.5s) with stronger NE activity compare to DA and 5HT (multiple comparisons, both p < 0.001). In the later activity (t = 17 to 20s), we found a significant neuromodulator main effect (F(2, 44) = 142.7, *p* < 0.0001) and significance of all three multiple comparisons (all *p* < 0.001). Similarly, in bout-related df/f, there was a significant neuromodulator main effect on both immediate (F(2, 44) = 44.08, *p* < 0.0001) and later activity (F(2, 44) = 9.842, *p* < 0.0001). Multiple comparisons further revealed a significantly stronger immediate response in DA versus NE (p < 0.01) and DA versus 5HT (p < 0.001), and NE versus 5HT (p < 0.001) in immediate activity, and both NE and DA were significantly stronger than 5HT (both p < 0.005) in later activity. Next, we aimed to determine whether the reward history would have an impact on dynamic changes by assessing the traces between the first 10 versus last 10 reward bouts. Both mPFC DA and NE showed an immediate peak at the first 10 bouts, and each trace was divergent from one in the last 10 bouts (**Figure S4A&B**). Collectively, these results suggested that NE plays a more prominent role representing negative valence, whereas DA encodes reward bout, and 5HT is negatively associated with both valences (**Figure 5A&S10**).

**Figure 5.**
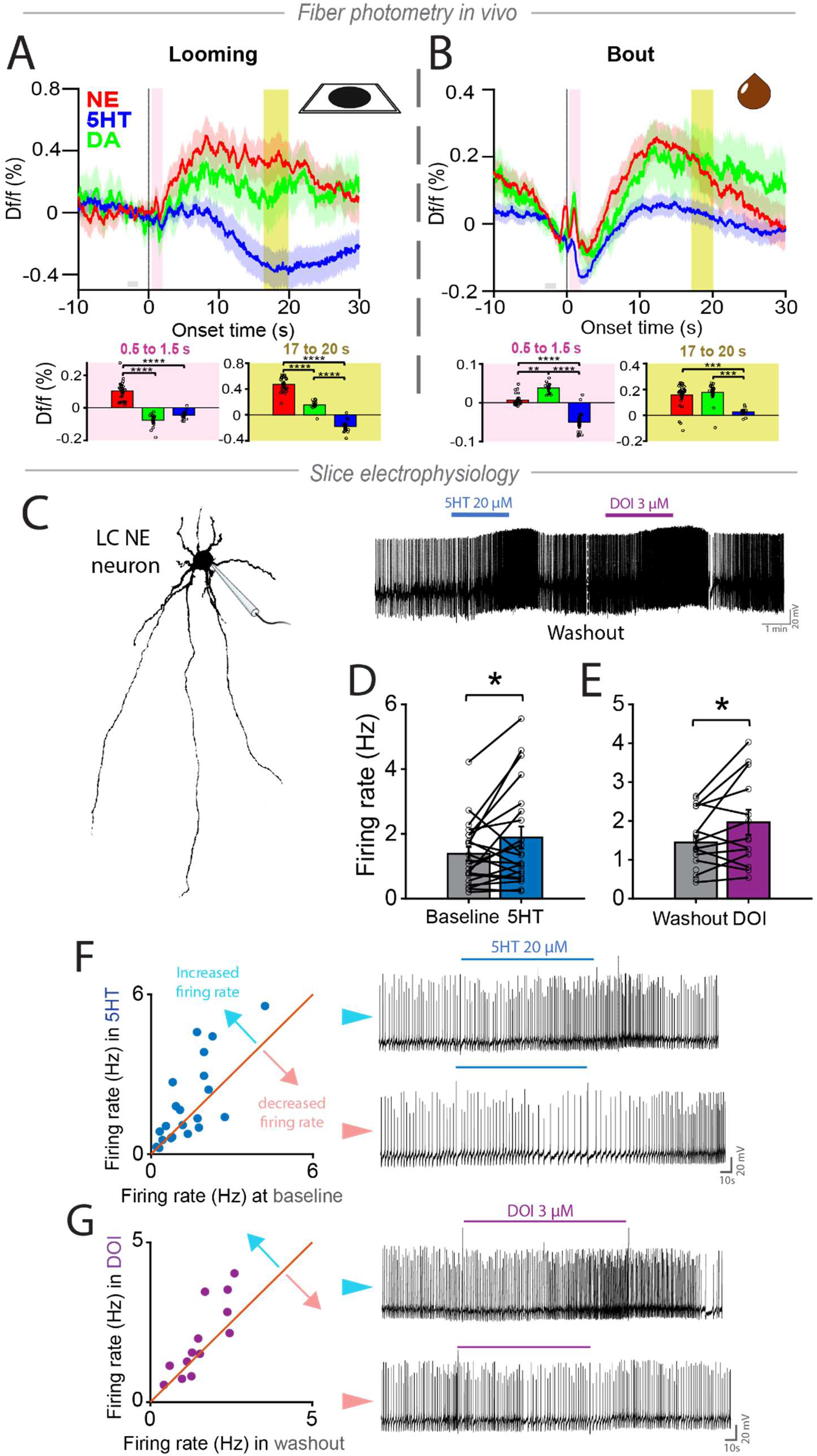
Interactions between 5HT and NE in vivo and in slice. **A-B.** NE, DA, and 5HT showed dissociative dynamics toward negative (i.e. looming threat) and positive (reward bout) valences. Looming-related (**A**) and bout-related (**B**) traces across all daily sessions. (Bottom) One-way ANOVA analyses of averaged activities (±S.E.M.) from 0.5 to 1.5s (pink) and 17 to 20s (mustard yellow) windows. **: p < 0.01; ***: p < 0.001; ****: p < 0.0001, multiple comparisons with adjusted alpha value. **C-G**. *Ex vivo* slice recording of locus coeruleus (LC) NE neurons with 5HT-related treatments. **C**: Diagram of patch clamp recording of an example LC NE cell body and its spontaneous activity. **D**: Spontaneous spikes of LC NE neurons after 5HT (20 µM), and **E**: a 5HT_2A/2C_ agonist DOI (3 µM) treatment. *: *p* < 0.05, paired t-test. **F:** Sub-population of LC NE neurons showed heterogeneous firing rate to 5HT (left). Example cells showing increased (top) and decreased (bottom) firing rate. **G:** Heterogeneous firing rate of LC NE neurons to DOI.

### 5HT heterogeneously modulates NE spiking

Since serotonin and norepinephrine have distinct neuromodulator release patterns in innate emotional behavior in this study, we reasoned that direct interactions from 5HT on LC NE could mediate their apparent emotional opponency. We conducted an *ex vivo* slice electrophysiology recording of LC NE neurons with 5HT-related treatments (**Figure 5C & S5**). Spontaneous spikes in LC NE neurons were significantly increased after bath-applied 5HT (*p* < 0.05) and a 5HT_2_ receptor agonist DOI (*p* < 0.05, **Figure 5D-E**). Notably, we found sub-populations of LC NE neurons showed no change or decreased firing rate to 5HT and DOI treatments (**Figure 5F-G; Figure S11**). To corroborate the functional relationship between these neurotransmitter systems, we performed in vivo fiber photometry of NE during looming while raising tonic NE or 5HT with systemic transporter inhibitors (**Figure S12**). We found that raising tonic NE reduced looming-induced NE release, while increasing 5HT did not alter looming-induced NE (**Figure S12**). Thus, these results support 5HT interactions with NE release being heterogeneous and receptor-dependent.

## Discussion

In this study, we developed a naturalistic approach avoidance task involving looming predation and reward and characterized prefrontal neuromodulatory dynamics during these motivational demands. Using fiber photometry of GRAB sensors, local pharmacology, and ex vivo slice electrophysiology, we report several findings. First, looming responses diminish over repeated days in this task and looming behavioral responses are diminished by silencing mPFC. Second, NE encodes responses to innate threat in this task showing selectivity for behavioral responses (higher with freezing) and across days (showing similar patterns of habituation). DA and NE both responded to innate threat and reward, but differed in their relative contributions with DA showing stronger reward responsivity. Local pharmacological antagonism of b-adrenergic receptors blocked freezing but not escape behaviors. Third, looming 5HT, in contrast showed suppression to looming exposure. Fourth, serotonergic modulation of LC firing may be heterogeneous based on slice electrophysiology data.

A major challenge in behavioral neuroscience is to design experimental paradigms capture the complexity of real-world decisions, particularly for emotional behaviors (Dennis et al., 2021; Zambetti et al., 2022; Miller et al., 2019). Approach-avoidance behaviors address a core construct in anxiety by juxtaposing the motivation to avoid threat and that to seek reward. Our task centers approach-avoidance around the ethological situation in which an animal has to leave a safe shelter to seek reward but also encounters the risk of experiencing predation (Yilmaz & Meister, 2013; Evans et al., 2018). The task intentionally used temporal unpredictability within the reward period to sustain the internal sense of threat. Over repeated sessions, animals displayed flexibility in their decision-making by reducing threat-driven behavioral responses while maintaining reward seeking. This adaptive behavior is crucial for survival because animals need to conserve energy from unnecessary escaping responses and exert effective exploitation of scarce resources (Evans et al., 2019; Dennis et al., 2021).

Prelimbic mPFC mediates both reward and aversive decisions and a thalamic-mPFC projection is involved in looming innate threat behavior (Salay et al 2018). Our finding that muscimol-induced suppression of mPFC diminishes innate threat behaviors further extends this work. Interestingly, a prior study found that cat predator exposure did not require prelimbic mPFC, perhaps suggesting a more refined stimulus specificity of innate threat circuitry (Corcoran and Quirk 2007). Cognitive circuits enable flexibility in the response to innate behaviors, underscoring their complexity (Leni et al 2022; Vale, Evans, Branco 2017). In a naturalistic context involving concurrent motivational drives, mPFC may be recruited to switch between affective goals even for innate behaviors. The observation that freezing was suppressed by b-adrenergic blockade and strongly drives prefrontal NE is consistent with a model in which noradrenergic signaling in mPFC stabilizes aversive states (Fitzgerald et al 2015) and guides defensive behavior strategies.

By measuring neuromodulatory dynamics with high resolution, we observed that each neuromodulator had a distinct pattern of encoding of motivational drives. Looming evoked time-locked responses in NE were stronger than those of DA and 5-HT. As the behavioral procedure was repeated over days, NE signals diminished along with the perceived salience of the threat; notably NE also responded to rewards. The common responsivity of DA to both reward and innate aversive stimuli could also be related to salience (Kutlu et al 2021). Biphasic responses were observed in PFC_NE_ in response to reward bout onset; a similar pattern was observed in LC dynamics during food consumption by Sciolino et al (2021) and could represent initial salience or orienting response followed by a subsequent suppression. DA and NE both encode a salience-like signal but there are important differences: DA shows greater responsivity to reward, while NE shows greater responsivity to threat. These observations may inform models in which neuromodulators may respond to both a common salience signal and show greater response towards specific stimuli. The fine-scale dynamics of NE also show transients at reward and during abrupt changes in state, consistent with some predictions of network-reset models (Sara and Bouret). Overall, these results may enhance integration of models of DA and NE, thus demonstrating the value of naturalistic tasks to elicit multidimensional emotional signals.

In contrast, 5HT showed decreased activity to both looming and reward bout. This finding is surprising because previous studies have suggested that DRN 5HT neurons are activated at reward intake and to reward-predicting cues (Liu et al, 2014; Cohen et al., 2015; Li et al., 2016; Matias et al., 2017; Ren et al., 2018). On the other hand, unlike reward signals, the results of DRN 5HT dynamic change to punishment are mixed. Studies reported DRN 5HT neurons have no response to aversive stimuli including timeout period, quinine, and foot shock (Liu et al., 2014; Li et al., 2016), but population-dependent excited or inhibited firing rate to air puff (Cohen et al., 2015). More recently, one study suggests that this punishment-evoked dissociation DRN 5HT is circuit-dependent, in which the central amygdala-projecting neurons were activated, whereas the orbitofrontal cortex-projecting neurons were inhibited, toward foot shock (Ren et al., 2018). We observed using slice electrophysiology that 5-HT regulation of LC neurons is receptor subtype–dependent and heterogeneous, in keeping with prior studies (Aghajanian et al., 1990; Aghajanian & Marek, 1999). Importantly, when we raised 5-HT tonically using fluoxetine, we did not observe any change in looming-evoked norepinephrine dynamics. Thus, opponent encoding of reward and innate threat between serotonin and norepinephrine may be mediated by other sources such as upstream circuits or may be elicited by other forms of 5-HT stimulation such as phasic release.

Integrating our findings, we propose a working model in which distinct streams of neuromodulatory information drive approach-avoidance decision-making in mPFC. Moment to moment changes in norepinephrine and dopamine may share common non-specific salience features to drive network resets or arousal shifts. At the same time, distinctive biases exist in which NE and DA demonstrate greater loadings onto innate threat and reward. Thus, NE and DA in the context of innate behaviors convey shared information about salience as well as having distinct biases towards a specific emotional valence. mPFC 5-HT may operate as an inhibitory signal towards innate valenced stimuli, in contrast to its distinctive effects on learned fear. This framework is broadly in agreement with prior theories of monoaminergic encoding elicited by Yu and Dayan (2005), while also extending to the distinctive features of innate approach-avoidance behavior.

There were several limitations of our study. Studying the activity of neurotransmitter dynamics in separate cohorts enabled comparisons between each neuromodulator and stimulus-driven responses; however, combining multiple neuromodulators would have provided additional insight into their interactions. While we provided causal tests of the role of mPFC and NE, phasic modulation of neuromodulatory signals could provide insights more closely related to the effects of neurotransmitter dynamics at the seconds scale. There have been multiple recent studies linking dynamics of one neuromodulator to direct effects on another locally; our study of neurotransmitter dynamics establishes the encoding differences during naturalistic behavior but does not address this distinct question. This point is particularly salient considering the heterogeneous interactions between 5-HT and LC, demonstrated by our results. Future studies combining phasic stimulation with neurotransmitter recordings in PFC (i.e. stimulation of 5-HT while measuring NE) may address this question more comprehensively in the context of ethologically relevant behaviors.

The present study characterized naturalistic dynamics of neuromodulators in mPFC as animals engaged in a task involving the presence of rewards and innate threats. Exaggerated threat perception or maladaptive decision-making about avoidance of danger are characteristic of anxiety and PTSD, and involve mPFC circuits. Understanding the neural circuit dynamics underpinning the switches between positive and negative valence behaviors may thus provide inside into the real time decisions that underlie mental health disorders. Important future directions include examining additional neuromodulators and neuropeptides in this paradigm (Zhu et al., 2018; Zhou et al., 2019; Barbano et al., 2020), employing projection-specific manipulations, and utilizing this task across developmental time points given the cross-species conservation of looming-evoked defenses (Schiff et al., 1962; Nanez, 1988; van der Weel & van der Meer, 2009). By grounding the study of neuromodulatory function in ethologically valid behavior under genuine motivational conflict, this approach may yield insights into adaptations to repeated traumatic experiences.

## Materials and Methods

### Animals

A total of 63 (male = 36, female = 27) adult C57BL/6J mice (Jackson Laboratory, 8 – 10 weeks old at arrival) were used in this study. Animals were housed in groups of up to four mice per cage on a 12/12-h light–dark cycle with lights off at 19:00. Water and food were provided *ad libitum* in home cages. All procedures were performed in accordance with the regulations of NIH guidelines and the Institutional Animal Care and Use Committee at Yale University.

### Apparatus

The behavior experiment was performed in a rectangular box (60 x 20 x 30 cm) built by clear acrylic sheets (adapted from Evans et al., 2018). A red-tinted acrylic shelter (10 x 19 x 13 cm) was placed at one end of the arena, and the reward spout (20 gauge blunt dispensing tips, CMLsupply) was inserted through a small drilled hole (4 cm from floor) on the wall at the other end. Each reward delivery was 3 µL of 50% Ensure (milk chocolate flavor, Abbott) mixed with water, and was dispensed via reward spout upon licks. Licks to the spout were detected by a custom made lickometer consisting of a dispensing tip electrically connected to a capacitive touch sensor and an Arduino redboard (Sparkfun Electronics). Reward delivery was controlled by scripts written in Presentation software (Neurobehavioral Systems, Inc.) via a DAQ card (USB-201, Measurement Computing) and a solenoid valve (McIntosh Controls Corp). A 27-inch monitor (1,920 x 1,080, Samsung SR35) was held facing down to the arena (60 cm from arena floor) to present visual looming stimuli. The looming stimulus was a video of a dark quick-expanding disk (full disk diameter 1,000 pixels) on white background generated by custom scripts written in Matlab (2021a, MathWorks, Inc.). A white noise generator (Homedics) set at 55-60 dB was used during behavioral experiments. Mouse behavior in each daily session was recorded by a camera (LifeCam HD-3000, Microsoft) mounted on the display monitor above the arena. The entire task structure—including lick detection and reward delivery—was automated and logged using custom scripts written in Presentation software.

### Approach/avoidance task

Mice were handled and pre-fed with 1 mL 50% Ensure solution in home cages for at least two days prior to behavioral experiments. Food and water were available in home cages throughout the study. The approach/avoidance task consisted of a one-day habituation and 5 approach/avoidance days. In each daily session, mice were firstly placed in a bedded empty cage with fiber cord connected to the optic fiber implant for 5 minutes to habituate the cables. Mice were then placed into the arena for 15 minutes without reward or looming stimulus on habituation day. On the following approach/avoidance days (25-30 min), reward was available upon licking the spout with a 250 ms inter-reward-interval due to the nature of rapid mouse licks (8-12 Hz). Looming onset was jittered within the reward period to preserve unpredictability of the innate aversive stimulus and thus to maintain sustained approach–avoidance conflict. If looming were presented at a predictable time relative to reward period onset, animals might adapt behavior to infer periods of safety rather than sustaining reward-danger conflict.. One looming stimulus consisted of 6-10 cycles of a quickly expanding dark disk on a white background (100% black-white contrast). Looming was conducted with average angular expansion rate of ∼15.8 deg/s (9.5° / 0.6 s); a lower angular speed compared with prior studies was used to provide an moderate aversive stimulus which could then be in conflict with appetitive drive.. The looming stimulus stayed at maximal for 400 ms, and followed by an 800 ms inter-cycle-interval. Looming stimuli were displayed only when the mouse was not in the shelter. The photometry system was calibrated and started when the cord-connected mouse was placed in the empty cage and throughout each daily session. Separate bedded empty cages were used between male and female mice. The arena and shelter were cleaned with 70% ethanol between mice.

The tone version of the approach/avoidance task consisted of one habituation day (same as described above) and 8 experimental days. On day 1 to 4 (training days, T1 to T4, 20 min), mice were placed in the arena with rewards available only when a 5k Hz tone was on (75-80 dB). Licks during tone-on periods were rewarded and recorded as ‘hits.’ In turn, licks during tone-off periods were not rewarded and were recorded as a “miss”. Tone-on durations (10, 15, …, 60s) and inter-tone-intervals (15, 16, …, 25s) were randomized with an averaged on/off ratio at around 63%. On day 5 to 8 (looming days, L1 to L4, 20 min), looming stimuli randomly presented twice a day only during tone-on periods. In tone-off periods, there was no reward or looming stimulus, and licks during which were recorded as ‘misses.’

### c-Fos whole brain mapping

Twelve C57BL/6 mice (6 males/6 females) of a new cohort were randomly assigned into a looming or novel cage control group in a new behavioral room (each group n = 6, 3 males/3 females). Mice in the looming group were placed in the approach/avoidance arena for 10 minutes with one looming stimulus (6-10 cycles) presented at 5th minute (**Figure S1**). Mice, including those in the control group, were then removed to an empty cage individually without food or water and rested for two hours. After which, mice were deeply anesthetized with isoflurane and perfused transcardially with PBS with 10U/mL heparin (Sigma-Aldrich), followed by ice-cold 4% paraformaldehyde (Sigma-Aldrich). Brains were collected and fixed in 4% paraformaldehyde at 4℃ for 24 hours with gentle shaking. The samples were then washed in PBS and stored in 0.02% sodium azide (Sigma-Aldrich) in PBS, and were shipped to LifeCanvas for whole brain c-Fos imaging. Positive cFos cells were identified by expressing both cFos (642 nm) and neuronal nuclei protein (561 nm). The dataset contains c-Fos expressing cell densities from 1,678 distinct brain areas with an imaging resolution of 1.8 x 1.8 x 4 microns per voxel. Left and right hemispheres of the same region were combined and averaged. Data were analyzed offline by customized scripts in Matlab.

### Surgery

#### Photometry

Mice underwent a surgery of viral injection and fiber implementation as described previously (Basu et al., bioRxiv). In brief, mice were anesthetized with 1-4% isoflurane (Covetrus, ME) in oxygen and secured in a stereotaxic frame. The scalp was first disinfected with sterile alcohol pads (Fisherbrand) and was partially removed to expose the dorsal aspect of the skull. The periosteum was gently removed, and the surface of the skull was cleaned with 0.9% saline. Bilateral injection sites were made by drilling through the skull at [AP] = +1.6 mm, [ML] = ±0.8 mm relative to bregma to expose the dura. Note the [ML] was adjusted from ±0.3 mm (above mPFC) to ±0.8 mm due to a 20° angle injection was applied to avoid the central sinus and to ensure space between two optic fibers (**Figure 2A&B**). For photometry experiments, 0.5 µL of GRAB sensors (AAV9-hSyn-GRAB sensor; either NE2h/DA3m/5HT2h, WZ Biosciences) and GRAB_DA_ (AAV9-hSyn-DA3m, WZ Biosciences) sensors were injected into mPFC unilaterally. Each virus was infused via a 5 µL Hamilton syringe (85N syringe, Hamilton, NV) targeted at depth [DV] = −1.4 mm with 20° angles. The injection rate was 0.1 µL per minute, and the syringe was left in place for an extra 5 minutes to minimize backflow of the injected solution. After viral injection, 3 mm optic fibers (Neurophotometrics, Ltd.) were implanted at depth [DV] = −1.4 mm from brain surface with 20° angles bilaterally and were secured by dental cement (C&B Metabond; Parkell Inc.) which also covered the entire exposed skull. After removal from surgical apparatus, mice were treated with lidocaine (5 mg/kg, Covetrus, ME) and carprofen (5 mg/kg, Butler Animal Health) subcutaneously. Post-operatively, carprofen and lidocaine treatments were applied daily for two days, and followed by a four-week grace period for viral expression before behavioral experiment commencement.

#### Infusion cannulas

Mice underwent the same surgery preparation as described above. Bilateral 26-gauge guide cannulas (Protech International, TX) targeting the medial prefrontal cortex (mPFC) were implanted at +1.6 AP, ±0.5 ML, −0.3 DV relative to dura, according to the mouse brain atlas (Paxinos & Franklin, 2019). Guide cannulas were secured to the skull with dental cement (C&B Metabond; Parkell Inc.), which also covered the remaining exposed skull surface. Dummy cannulas were inserted into each guide cannula to maintain patency, and protective dust caps were secured over the cannula assembly. Internal cannulas (33-gauge) projected 1 mm beyond the guide cannula tip to reach the target region. Mice were allowed to recover for one week before behavioral testing.

#### Histology

After all experiments, mice were first deeply anesthetized with isoflurane and were perfused transcardially with ice-cold PBS followed by 4% paraformaldehyde (Sigma-Aldrich). Brains were carefully removed and post-fixed in 4% paraformaldehyde at 4℃ for at least 24 hours, and then switched to 30% sucrose solution until brain slicing. Frontal areas of the brains were sectioned coronally into 100 μm slices with a cryostat (Leica CM3050S) and mounted onto charged slides. Slides were covered by cover glass (Gold Seal, Thermo Scientific) with a mountant with DAPI (ProLong, Thermo Scientific), and then were imaged using a confocal microscope (Olympus FV3000) for fluorescence expression. The brightness and contrast of the images were adjusted appropriately in ImageJ to enhance the clarity of the area. Based on the histology, mice were excluded from analysis if fluorescence was not visible in mPFC region or a dislocation of fiber track from fluorescent areas.

#### Fiber photometry recording

The same fiber photometry system was also used to measure the GRAB sensor fluorescence. Fiber recording was achieved by connecting parallel patch cords to bilateral optic fiber implants in mice, enabling readout of sensor signals in mPFC. In each recording channel, light at the 470 nm excitation wavelength and a 415 nm isosbestic control was emitted from LEDs with 10% power and 20 Hz pulse frequency, and propagated to the optic fiber implants. Fluorescence emission from GRAB sensors was captured by an sCMOS camera at a frame rate of 40 Hz. The time-dependent fluorescence data were saved and analyzed offline by customized Matlab scripts.

#### Intracranial Infusion

Prior to infusion, the dummy cannula was removed from the guide cannula and an injection cannula was inserted. Mice were infused with either saline, 5 µg propranolol, or 0.5 µg muscimol. Drug order was counterbalanced within each experiment, with a two-day washout period between drug and saline sessions. Propranolol and muscimol experiments were conducted two weeks apart, each with their own saline control sessions. Mice were infused in a bedded, empty cage at a rate of 0.08 µl/min for a total volume of 0.2 µl per hemisphere. Following completion of the infusion, the injection cannula was left in place for 1 minute to allow for diffusion and prevent backflow, after which it was removed and the dummy cannula replaced. Mice remained in the cage for an additional 10 minutes before being transferred to the arena for the approach-avoidance task.

### Data Analyses

#### Behavioral data

All behavioral raw data including timestamps for licks, reward deliveries, and looming events were logged into text files by Presentation in each daily session. Data were pre-processed and analyzed by customized scripts written in Matlab. One reward bout was defined as a train of licks within 5 seconds from the last lick. Looming events were aligned by the start of looming stimulus and reward bout events were aligned at the first lick of each bout (onset time, t = 0). Looming-induced escaping behavior was defined as mouse running and reached to shelter within the first 3 cycles of looming stimulus, whereas freezing behavior was defined as rigidness without any visible movements. In contrast, ‘no response’ to looming was defined as behaviors other than freezing and escaping, including but not limited to grooming, exploring, or licking. Additionally, mouse behavior was recorded throughout each daily session and analyzed by a supervised tracking system (Deeplabcut, Mathis et al., 2018) for precise position tracking per frame. The model was trained by feeding ∼1,000 randomly selected frames, in which mice were manually labeled with the nose, left ear, right ear, central body, left side of body, right side of body, tail base, and tail end on the mouse (**Figure S2A**). The outputs generated by the well-trained network were analyzed by customized Matlab scripts.

#### Photometry data

GRAB sensor fluorescent signals were recorded and saved by Bonsai interface, and analyzed by customized scripts written in Matlab. In addition to histology examinations, certain sessions were excluded due to motion artifacts caused by twisted patch cords during recording. For data processing, the 470 nm traces were firstly smoothed with a non-linear regression algorithm to correct the mild photobleaching effect in each day. To assess the event-related dynamics, the fluorescent signals were extracted with certain time windows aligned with event onset time (t = 0). The event-related fluorescence Δ*F/F(t)* was then calculated for each time bin *t* as:

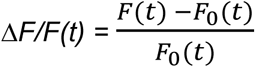

where *F(t)* is the fluorescence in time bin (50 ms each), and *F*_O_(*t*) is the baseline average signal ranging from −3 to −1.5 seconds from each event onset. To examine the day-by-day effects, all event-related signals in each session were averaged and plotted with 95% confidence intervals.

#### Ex vivo slice physiology

Brain slice preparation and electrophysiology recording procedures were similar to that described previously (Liu & Aghajanian, 2008). In brief, four C57BL/6 mice were used and their brains were placed in −4°C artificial cerebrospinal fluid (ACSF) in which sucrose (252 mM) replaced NaCl (sucrose-ACSF) to prevent cell swelling. Next, the brain tissue containing LC was sliced with 400 µm thickness using a vibratome (Leica VT1000S). Testing drugs were dissolved in ACSF and applied with a flow rate at 4 mL/min. A recovery period of ∼1–2 hr was allowed before commencement of recording. To visualize the NE neurons in LC, an Olympus BX50WI microscope (x40 or x60 IR lens) with infrared differential interference contrast (IR/DIC) videomicroscopy (Olympus) was used. Whole-cell recordings were performed with an Axoclamp-2B amplifier (Axon Instruments). The output signal was low-pass-filtered at 3 KHz, amplified x100 through Cyberamp, digitized at 15 kHz, and acquired by using pClamp 10.1/Digidata 1320 software (Axon Instruments). Postsynaptic currents were studied in the continuous single-electrode voltage-clamp mode (3-kHz low-pass filter cutoff frequency). After completion of recording, slices were transferred to 4% paraformaldehyde in 0.1 M phosphate buffer and stored overnight at 4°C. Slices were then processed with streptavidin conjugated to Alexa 594 (1/1,000; Invitrogen) for visualization of Neurobiotin-labeled cells.

## Statistics

All statistics were performed in Matlab or GraphPad Prism (9.4.1). No statistical analysis was employed to determine sample sizes; however, they were similar to those used elsewhere in the field. For behavioral data statistical analyses, one-way analysis of variance (ANOVA) with repeated measures was used to analyze the behavioral parameters. Post-hoc analyses with adjusted alpha value were performed to assess the differences in certain time windows if the main effect was significant. Fluorescence(df/f) was calculated relative to time-locked events with 95% confidence intervals obtained by resampling (bootstrap; 1,000 iterations). Chi-square test and student’s paired t-test were used to determine differences between treatments in slice physiology experiments. All data were presented as mean ± SEM or mean ± 95% confidence interval, and significance level was set at α = 0.05 unless otherwise noted. Data plots were generated by Matlab or Prism, and diagrams were made with Adobe Illustrator (27.1.1) and Biorender (BioRender.com).

## Data availability

The behavioral and photometry data, as well as customized codes written in Matlab and Presentation used in this study are available from the corresponding authors upon request.

---

**Figure S1.**
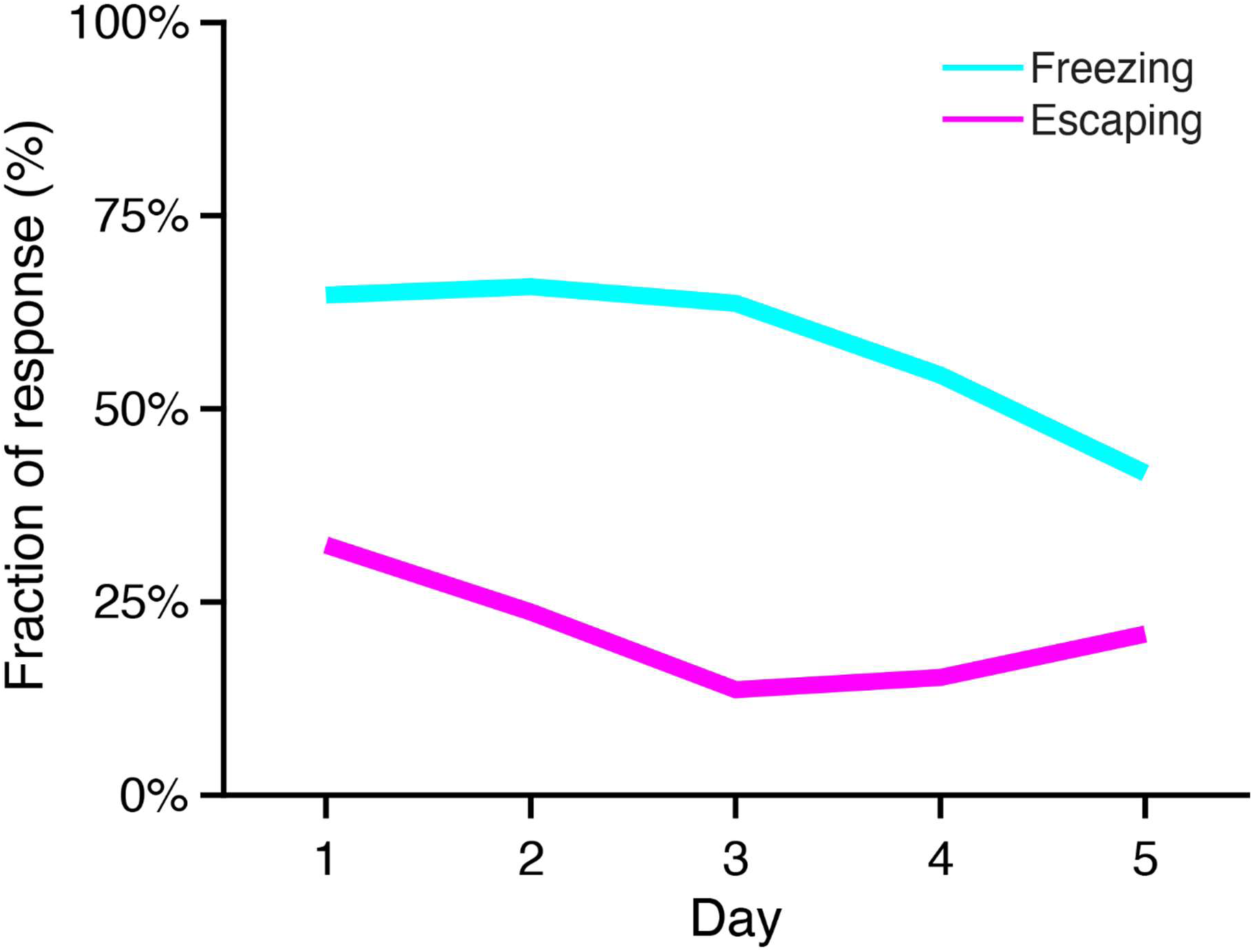
Distribution of fear behavior over days. Freezing and escaping over days.

**Figure S2.**
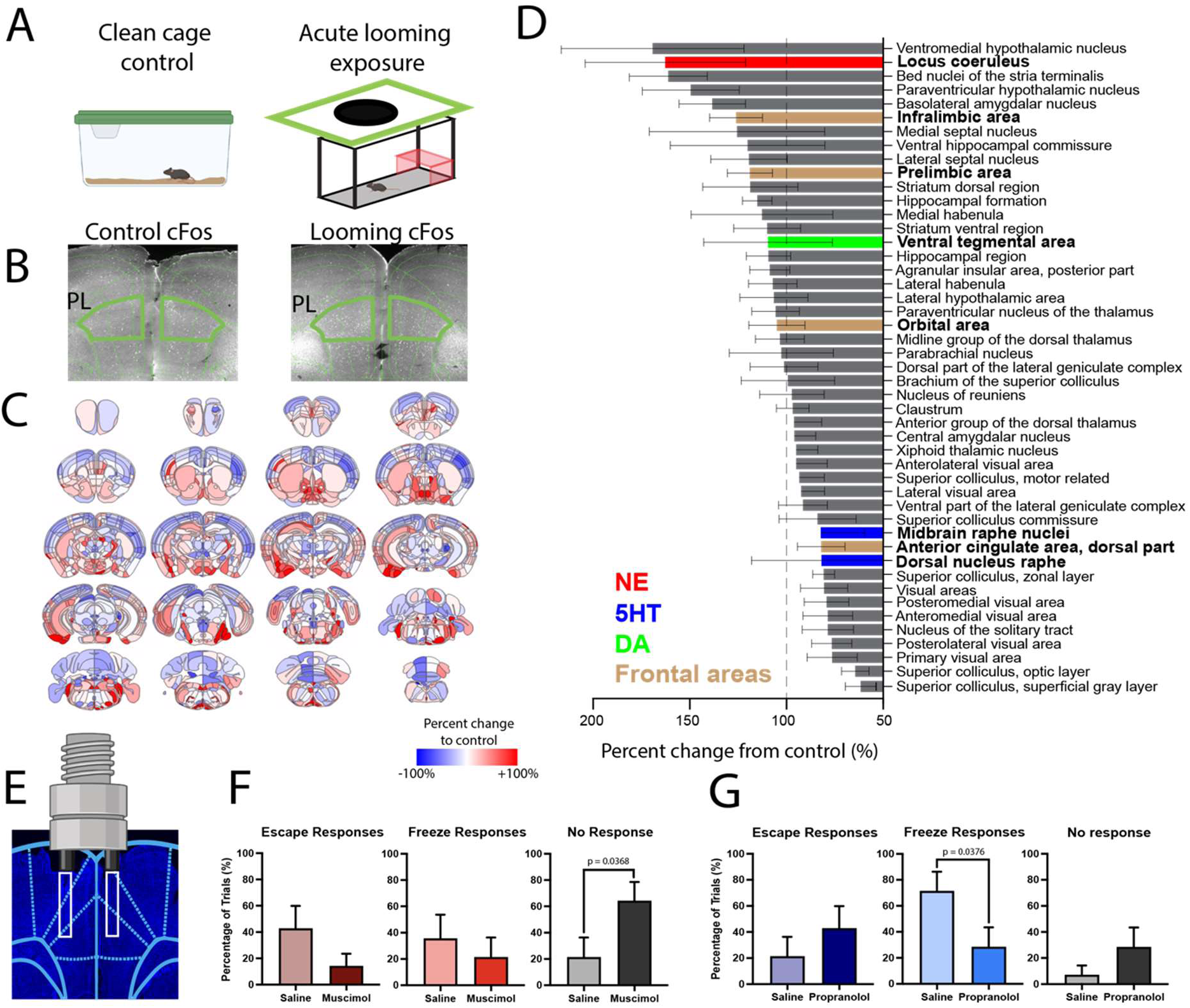
Whole-brain cFos mapping. **A**: Samples were collected from mice that experienced looming stimuli (left, n = 6), and baselined with a control group exposed to a new cage in a novel context (right, n = 6). **B**: Exemplar cFos staining images of PL in mice that either underwent novel cage exposure (control, left) or acute looming exposure (right). **C**: Averaged cFos expression in example coronal slices is shown in percent change of density from control (±S.E.M.). **D**: A list of regions of interest relating to looming stimulus. **E**: Example histology photo from mouse implanted with microinfusion cannula in mPFC. White boxes indicate internal cannula depth. **F & G**: Percentage of looming trials that resulted in either escape (left panel), freezing (middle panel), or no response (right panel) in animals treated with either saline or muscimol (**F**) and saline or propranolol (**G**).

**Figure S3.**
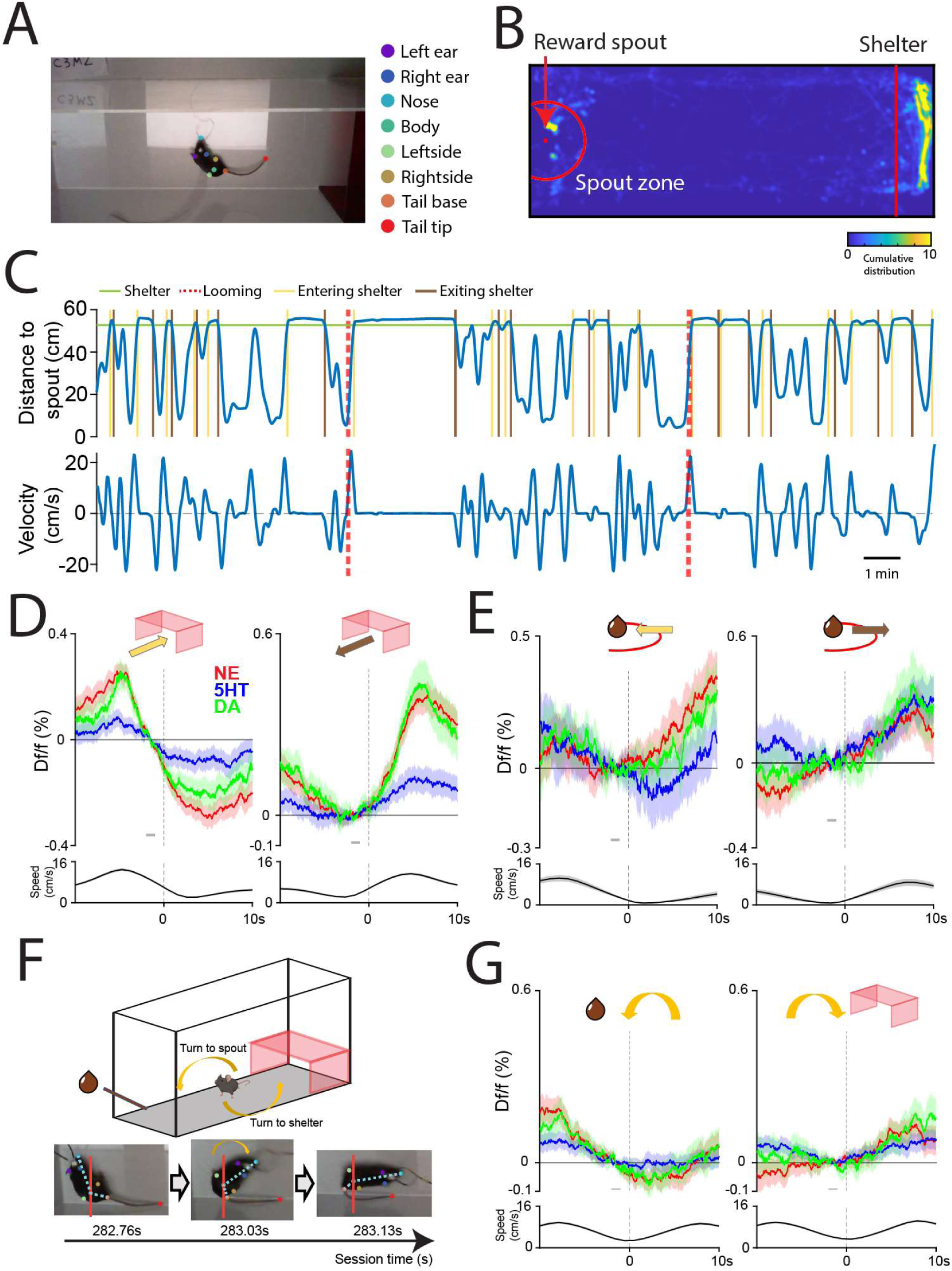
Behavioral events tracking by using a supervised deep learning model (Deeplabcut). **A**: An example frame of a mouse performing the task with labeled body parts. **B**: Heatmap of mouse position in cumulative distribution. The red dot represents the reward spout location, and the red circle represents the spout zone, which is defined as a radius of 10% of box length. And the red line represents the entering edge of the shelter. **C**: Example trace of a mouse traveling in the box in a daily session. Mouse position is shown by distance to spout (top) and running velocity (bottom). **D**: Neuromodulator activity at events of entering (left) and exiting (right) the shelter. Bottom panels represent the running speed. Shaded area indicates the 95% of confidence intervals. **E**: Neuromodulator activity at events of entering (left) and exiting (right) spout zone. **F**: A diagram illustrates turning to spout versus turning to shelter. (Bottom) Example frames of a mouse making a turn. **G**: Neuromodulator activity at events of turn to spout (left) and turn to shelter (right) spout zone.

**Figure S4.**
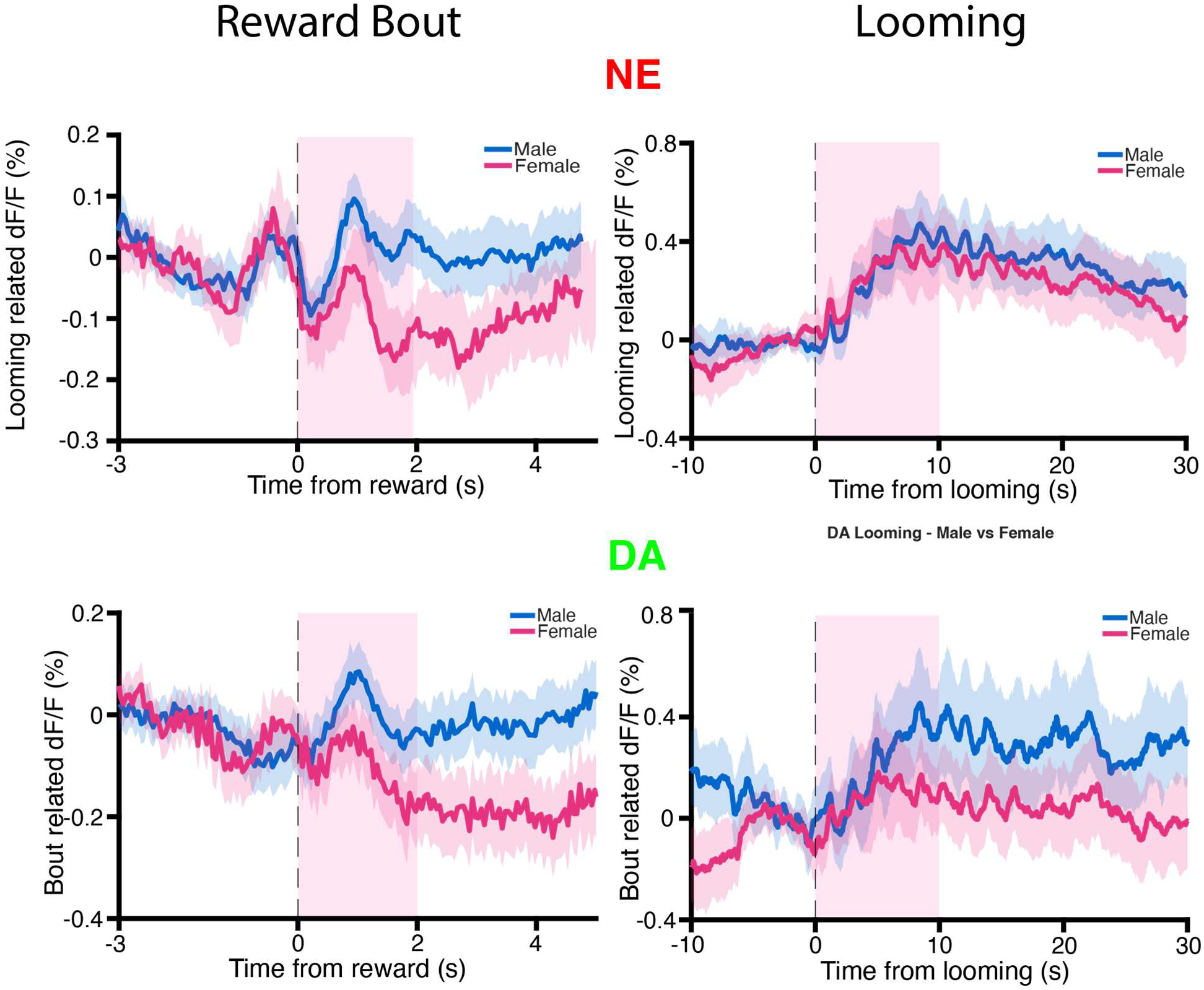
Neuromodulator dynamics in conflict task in male and female mice. (A) Average number of licks per session in males and females. (B–C) Average ΔF/F in response to reward bout (left) and looming stimulus (right) for NE (B) and DA (C). NE reward bout (0–2 s): LME, β = 0.098, t(1234) = 3.14, p = 0.002; looming (0–10 s): LME, β = 0.0004, t(204) = 0.34, p = 0.73, n.s. DA reward bout (0–2 s): LME, β = 0.092, t(523) = 2.11, p = 0.035; looming (0–10 s): LME, β = 0.0017, t(98) = 1.08, p = 0.28, n.s.

**Figure S5.**
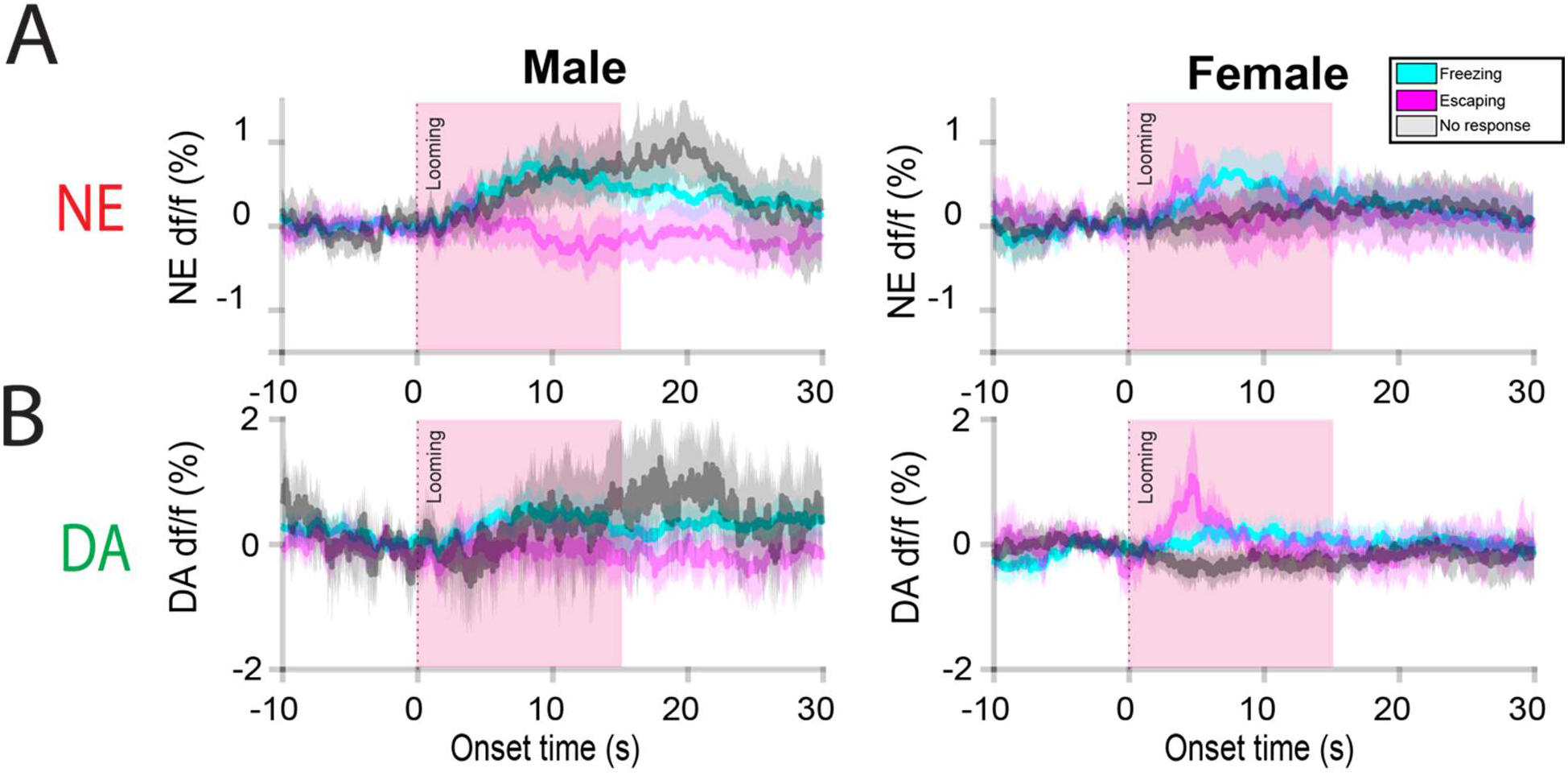
Neuromodulator dynamics during fear response behaviors in male and female mice. (A) Norepinephrine ΔF/F during freezing, escaping, and no response following looming stimulus presentation in males (left) and females (right). (B) Dopamine ΔF/F during the same behavioral responses in males (left) and females (right).

**Figure S6.**
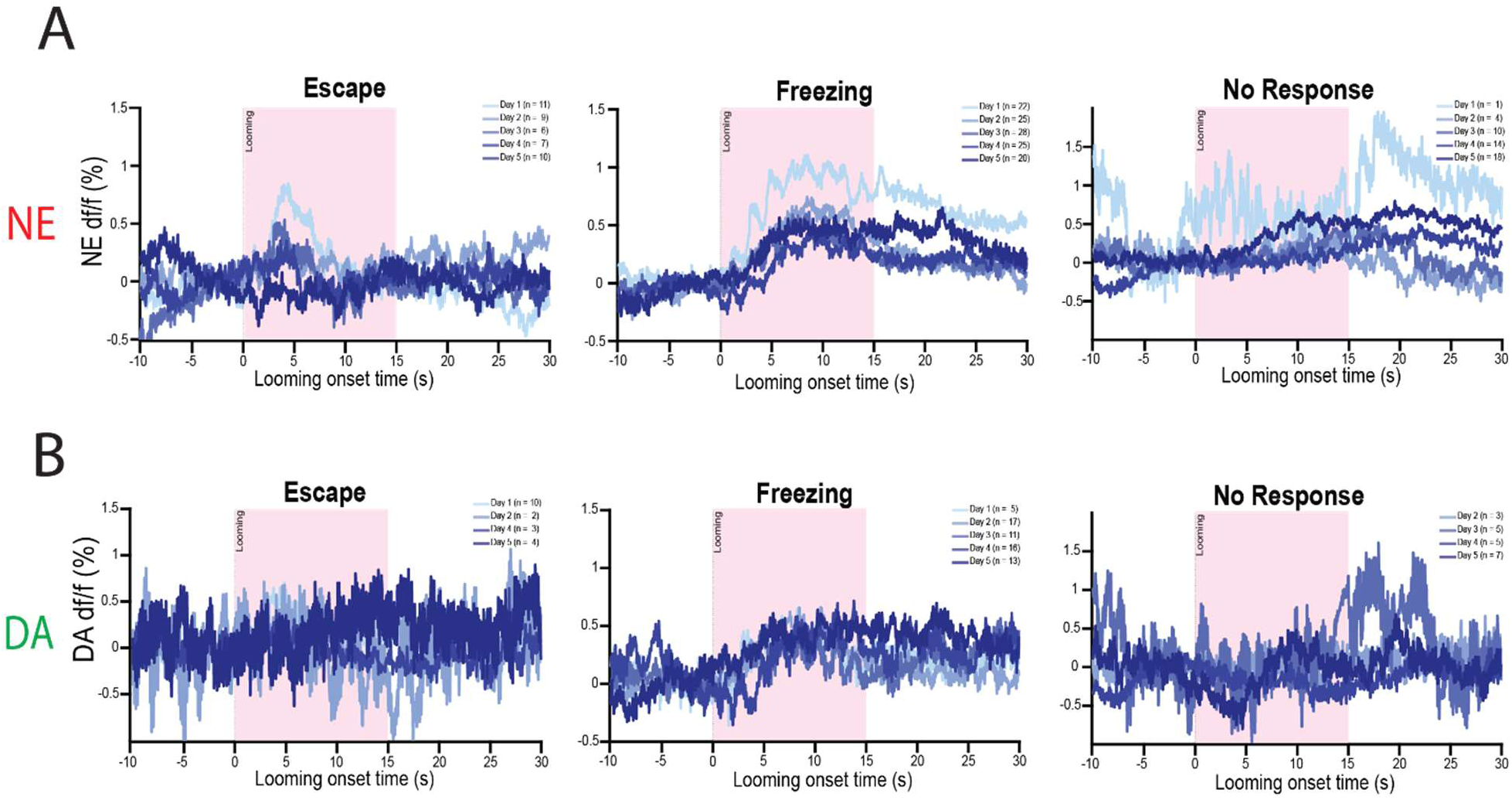
Neuromodulator dynamics during fear response behaviors across days. (A) Norepinephrine ΔF/F during escape (left), freezing (middle), and no response (right) following looming stimulus presentation across sessions. (B) Dopamine ΔF/F during the same behavioral responses across sessions.

**Figure S7.**
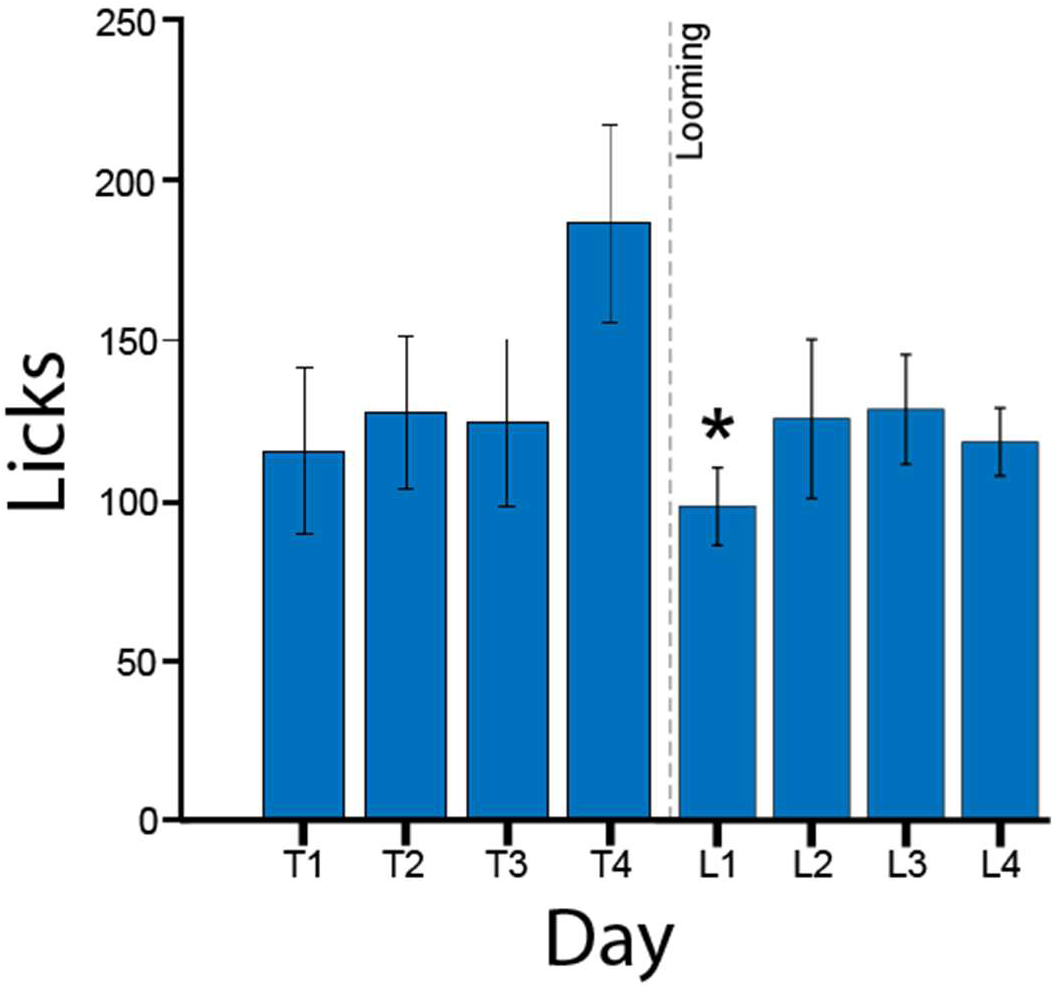
Addition of looming stimulus temporarily decreases average number of licks. Average licks over course of 4 training days and 4 conflict days. Licking was significantly decreased on day L1 in comparison to T4 (p <0.05, planned comparisons).

**Figure S8.**
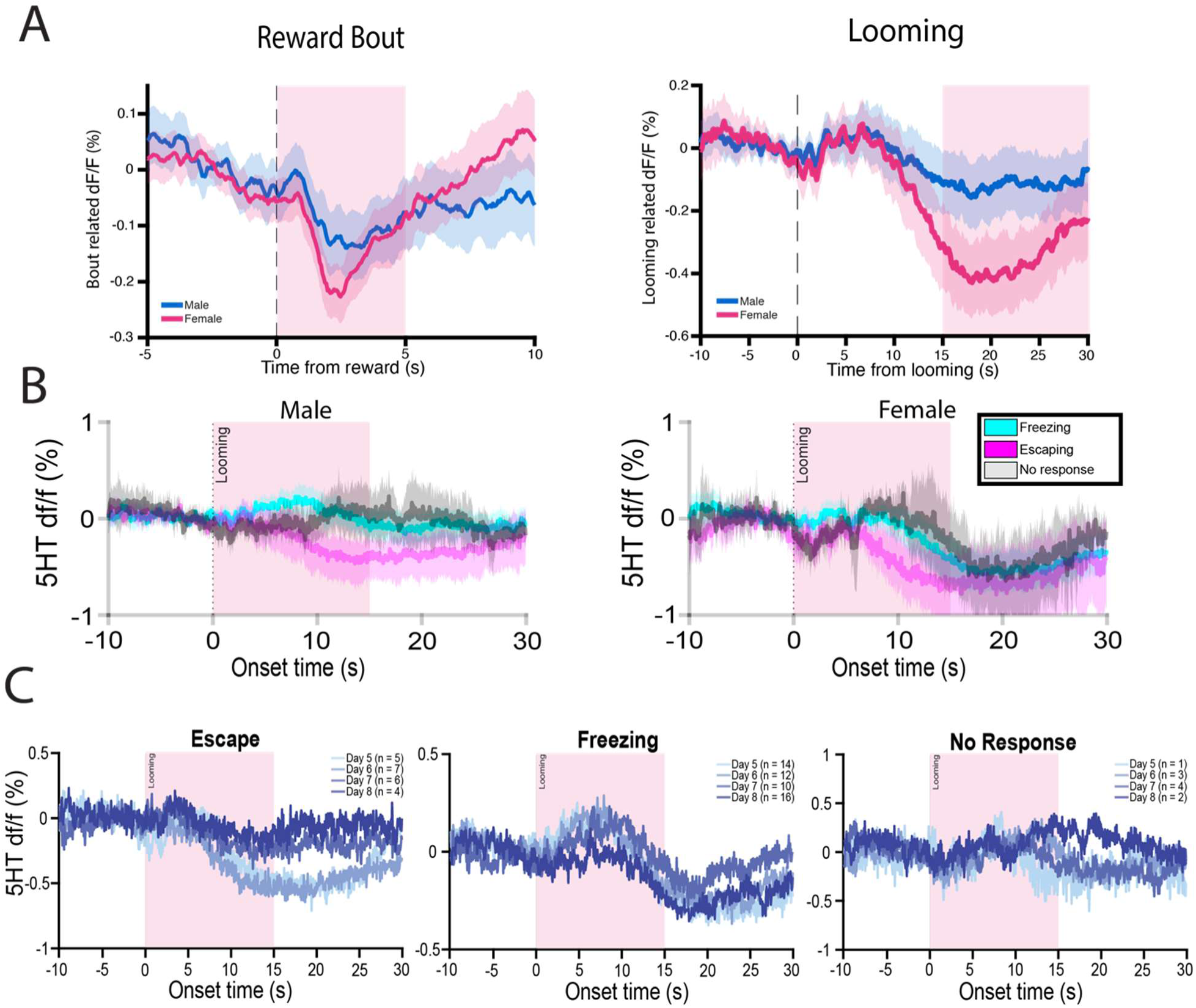
Serotonin dynamics during conflict task. (A) 5-HT ΔF/F in male and female mice during reward bout (left) and looming stimulus (right). Reward bout (0–5 s): LME, β = 0.041, t(388) = 1.46, p = 0.14, n.s.; looming (15–30 s): LME, β = 0.0023, t(82) = 1.90, p = 0.061, n.s. (B). 5-HT ΔF/F during freezing, escaping, and no response following looming stimulus presentation in males (left) and females (right). (C) 5-HT ΔF/F during escape (left), freezing (middle), and no response (right) across sessions.

**Figure S9.**
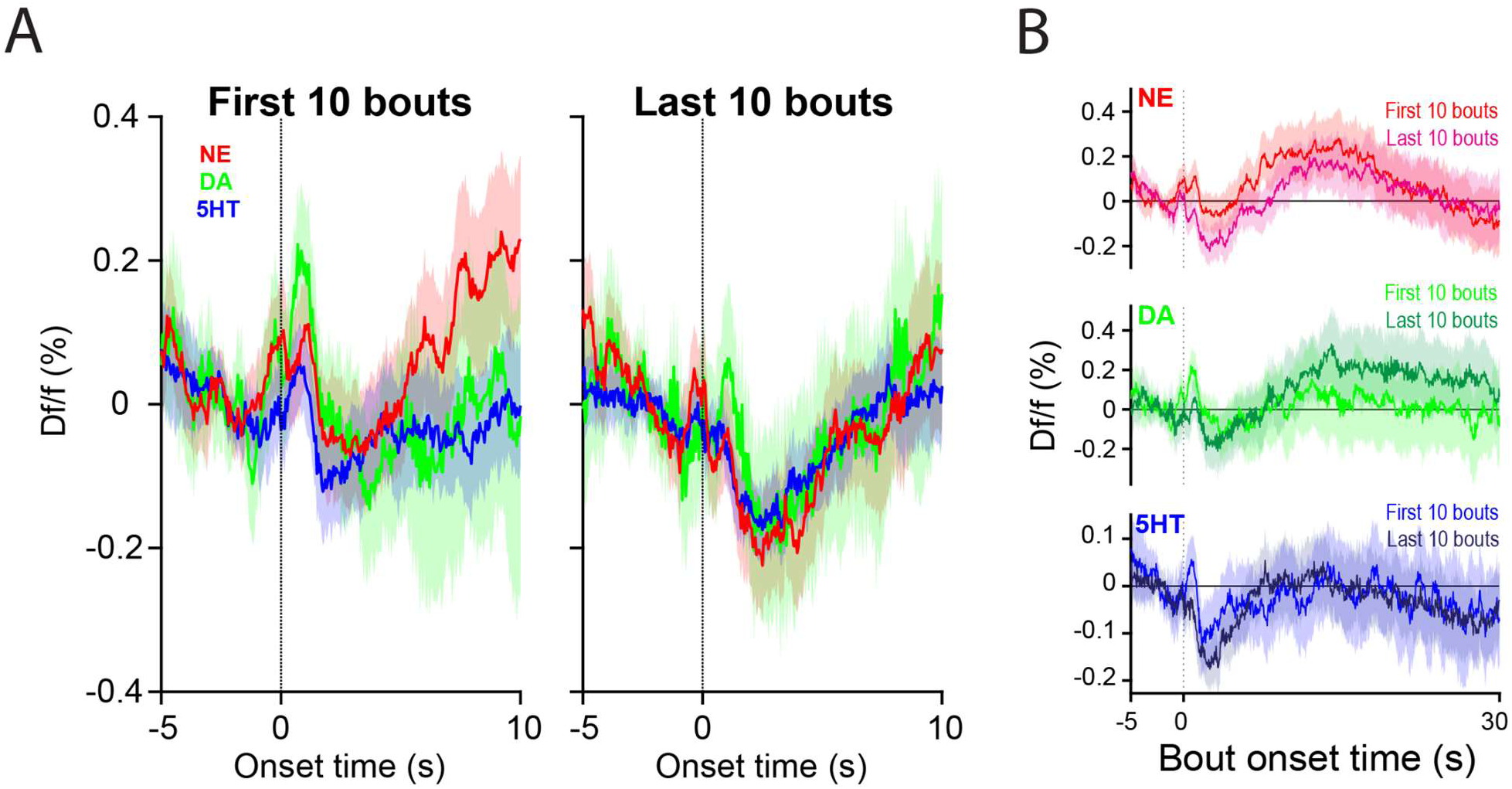
Neuromodulator activities respond to different bout states. **A**: Averaged NE, DA, and 5HT associated with the first 10 (left) and the last 10 bout (right). **B**: Visualization of the first versus last 10 bouts in NE (top), DA (middle), and 5HT (bottom) dynamics. Shaded area indicates the 95% of confidence intervals.

**Figure S10.**
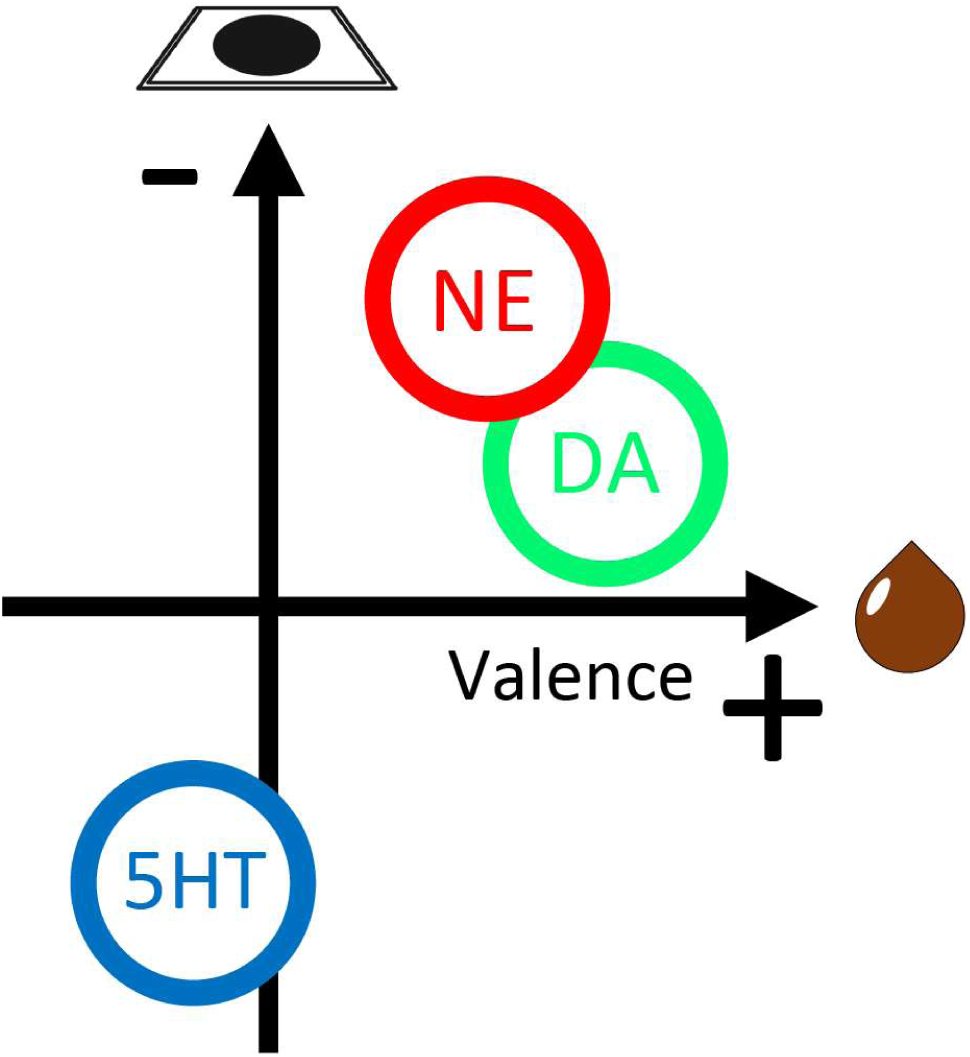
Schema of the roles of NE, DA, and 5HT in negative and positive valences. NE plays a more prominent role in representing threat (negative valence) and DA encodes reward (positive valence). In contrast, 5HT shows decreased activity to both valences.

**Figure S11.**
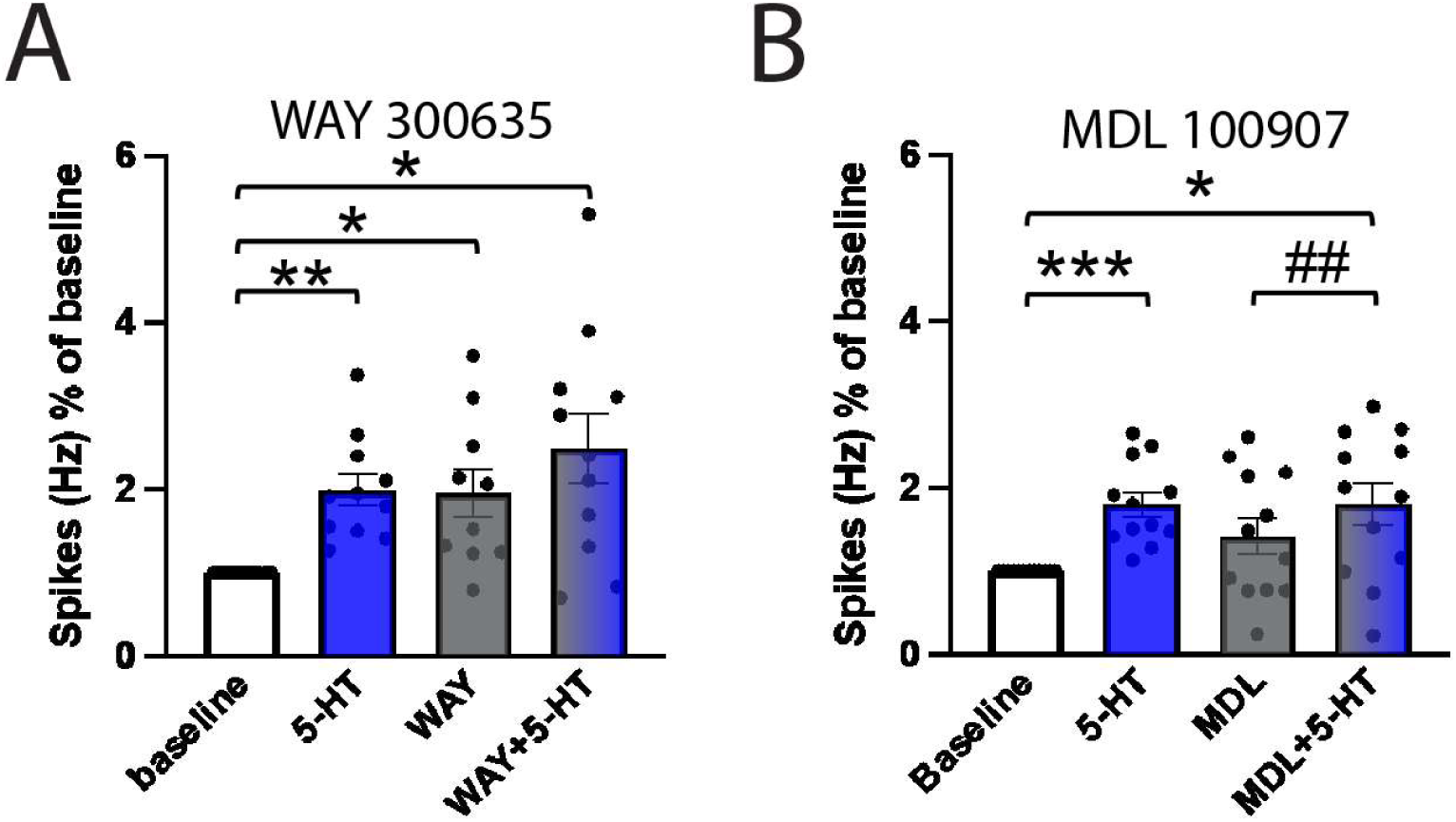
Effects of 5HT-related bath applications on LC NE neuron activity *ex vivo*. **A**: Effects of WAY300635, a 5HT_1A_ antagonist, on LC NE neurons spikes. **B**: Effects of MDL100907, a 5HT_2A_ antagonist, on LC NE neurons spikes. *: *p* < 0.05; **: *p* < 0.01; ***: *p* < 0.005, chi-square test. ##: *p* < 0.01, paired t-test.

**Figure S12.**
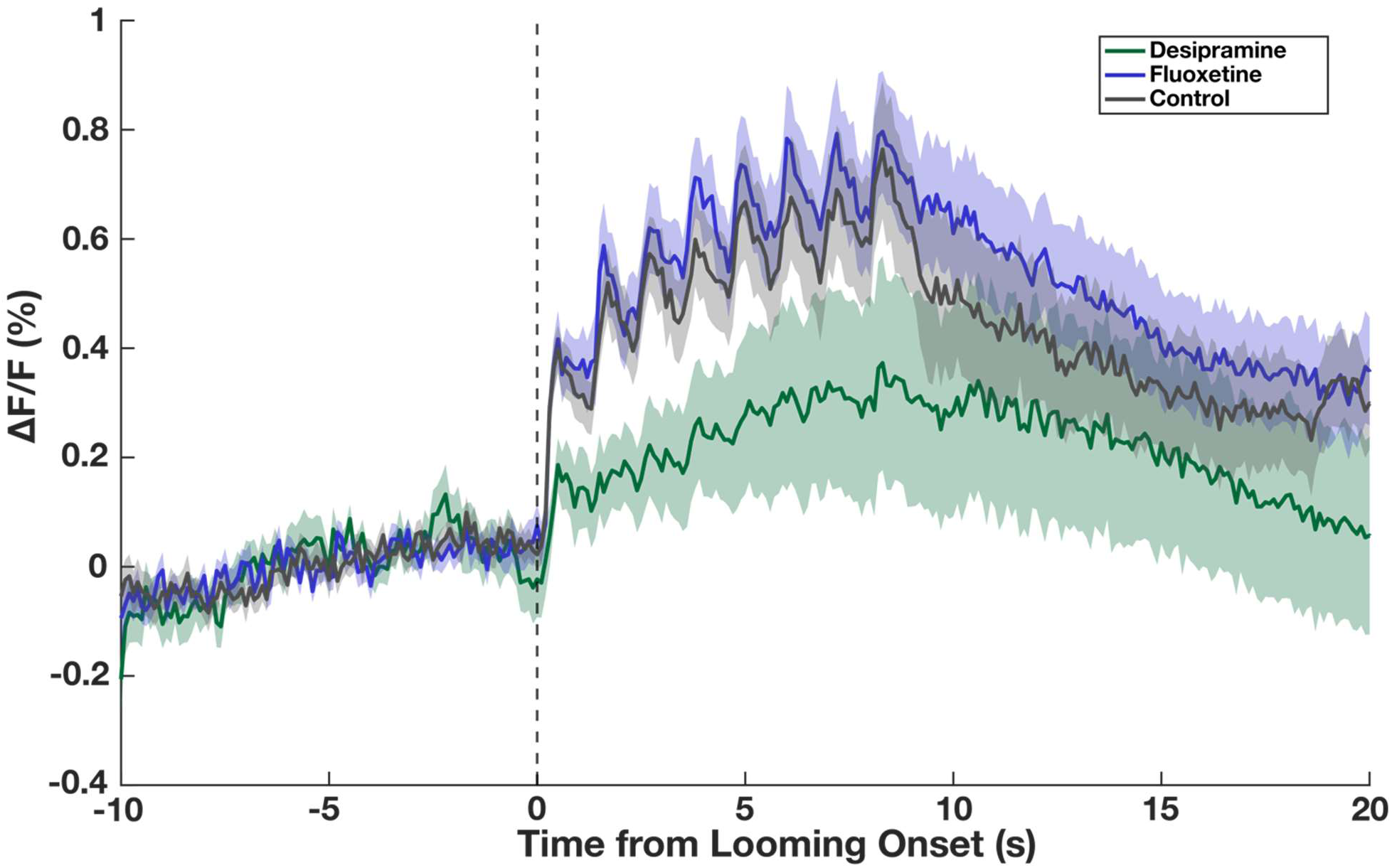
Looming-evoked NE ΔF/F is altered by increased NE but not by increased 5-HT. NE dynamics during looming stimulus presentation in mice pretreated with saline (control), desipramine, or fluoxetine. Shaded areas are bootstrap 95% C.I.. Mice treated with desipramine showed slightly decreased looming-evoked NE (RM-ANOVA: F(2,24) = 6.92, p = 0.004; Des vs Control post-hoc: p = 0.058)

## Notes

### Competing Interest Statement

A.P.K received research funding from Freedom Biosciences and Transcend Therapeutics. A.P.K., A.L.Y. submitted a patent application unrelated to this work.

### Summary of Updates

Added additional local pharmacology; added co-first author Emily L Burke

## References

Adolphs R. (2013). The biology of fear. Curr Biol, 23(2):R79–93

Aghajanian GK, Marek GJ. (1999). Serotonin and hallucinogens. Neuropsychopharmacology, 21(2 Suppl):16S–23S

Aghajanian GK, Sprouse JS, Sheldon P, Rasmussen K. (1990). Electrophysiology of the central serotonin system: receptor subtypes and transducer mechanisms. Ann N Y Acad Sci, 600:93–103

Aston-Jones G, Akaoka H, Charléty P, Chouvet G. (1991). Serotonin selectively attenuates glutamate-evoked activation of noradrenergic locus coeruleus neurons. J Neurosci, 11(3):760–9

Barbano MF, Wang HL, Zhang S, Miranda-Barrientos J, Estrin DJ, Figueroa-González A, Liu B, Barker DJ, Morales M. (2020). VTA Glutamatergic neurons mediate innate defensive behaviors. Neuron, 107(2):368–382

Basu A, Yang JH, Yu A, Glaeser-Khan S, Feng J, Krystal JH, Li Y, Kaye AP. (2022). Prefrontal norepinephrine represents a threat prediction error under uncertainty. *bioRxiv*, 2022.10.13.511463

Bedoya-Pérez MA, Smith KL, Kevin RC, Luo JL, Crowther MS, McGregor IS (2019). Parameters That Affect Fear Responses in Rodents and How to Use Them for Management. Frontiers in Ecology and Evolution, 7:136

Burnett CJ, Li C, Webber E, Tsaousidou E, Xue SY, Brüning JC, Krashes MJ. (2016). Hunger-Driven Motivational State Competition. Neuron, 92(1):187–201

Chandler DJ, Gao WJ, Waterhouse BD. (2014). Heterogeneous organization of the locus coeruleus projections to prefrontal and motor cortices. Proc Natl Acad Sci USA, 111(18):6816–21

Chen JA, Li S, Wang BX, Wu N, Li F, Li J. (2022). The effect of visually evoked innate fear on reward-associated conditional response and reversal learning in mice. Physiol Behav, 244:113648

Clement HW, Gemsa D, Wesemann W. (1992). Serotonin-norepinephrine interactions: a voltammetric study on the effect of serotonin receptor stimulation followed in the N. raphe dorsalis and the Locus coeruleus of the rat. J Neural Transm Gen Sect, 88(1):11–23

Cohen JY, Amoroso MW, Uchida N. (2015). Serotonergic neurons signal reward and punishment on multiple timescales. Elife, 4:e06346

Cools R, Arnsten AFT. (2022). Neuromodulation of prefrontal cortex cognitive function in primates: the powerful roles of monoamines and acetylcholine. Neuropsychopharmacology, 47(1):309–328

Delcourte S, Etievant A, Haddjeri N. (2021). Role of central serotonin and noradrenaline interactions in the antidepressants’ action: Electrophysiological and neurochemical evidence. Prog Brain Res, 259:7–81

Dennis EJ, El Hady A, Michaiel A, Clemens A, Tervo DRG, Voigts J, Datta SR. (2020). Systems neuroscience of natural behaviors in rodents. J Neurosci, 41(5):911–919

Evans DA, Stempel AV, Vale R, Branco T. (2019). Cognitive control of escape behaviour. Trends Cogn Sci, 23(4):334–348

Evans DA, Stempel AV, Vale R, Ruehle S, Lefler Y, Branco T. (2018). A synaptic threshold mechanism for computing escape decisions. Nature, 558(7711):590–594

Feng J, Zhang C, Lischinsky JE, Jing M, Zhou J, Wang H, Zhang Y, Dong A, Wu Z, Wu H, Chen W, Zhang P, Zou J, Hires SA, Zhu JJ, Cui G, Lin D, Du J, Li Y. (2019). A genetically encoded fluorescent sensor for rapid and specific in vivo detection of norepinephrine. Neuron, 102(4):745–761.e8

Fernández-Pastor B, Ortega JE, Meana JJ. (2013). Involvement of serotonin 5-HT3 receptors in the modulation of noradrenergic transmission by serotonin reuptake inhibitors: a microdialysis study in rat brain. Psychopharmacology (Berl), 229(2):331–44

Florin-Lechner SM, Druhan JP, Aston-Jones G, Valentino RJ. (1996). Enhanced norepinephrine release in prefrontal cortex with burst stimulation of the locus coeruleus. Brain Res, 742(1-2):89–97

Fratzl A, Koltchev AM, Vissers N, Tan YL, Marques-Smith A, Stempel AV, Branco T, Hofer SB. (2021). Flexible inhibitory control of visually evoked defensive behavior by the ventral lateral geniculate nucleus. Neuron,109(23):3810–3822

Ganella DE, Kim JH. (2014). Developmental rodent models of fear and anxiety: from neurobiology to pharmacology. Br J Pharmacol, 171(20):4556–74

Giustino TF, Maren S. (2018). Noradrenergic modulation of fear conditioning and extinction. Front Behav Neurosci, 12:43

Hu Y, Chen Z, Huang L, Xi Y, Li B, Wang H, Yan J, Lee TMC, Tao Q, So KF, Ren C. (2017). A translational study on looming-evoked defensive response and the underlying subcortical pathway in autism. Sci Rep, 7(1):14755

Huang L, Yuan T, Tan M, Xi Y, Hu Y, Tao Q, Zhao Z, Zheng J, Han Y, Xu F, Luo M, Sollars PJ, Pu M, Pickard GE, So KF, Ren C. (2017). A retinoraphe projection regulates serotonergic activity and looming-evoked defensive behaviour. Nat Commun, 8:14908

Hwang A, Skarica M, Xu S, Coudriet J, Lee CY, Lin L, Terwilliger R, Sliby AN, Wang J, Nguyen T, Li H, Wu M, Dai Y, Duan Z, Srinivasan SS, Zhang X, Lin Y, Cruz D, Deans PJM; Traumatic Stress Brain Research Group; Huber BR, Levey D, Glausier JR, Lewis DA, Gelernter J, Holtzheimer PE, Friedman MJ, Gerstein M, Sestan N, Brennand KJ, Xu K, Zhao H, Krystal JH, Young KA, Williamson DE, Che A, Zhang J, Girgenti MJ. (2025). Single-cell transcriptomic and chromatin dynamics of the human brain in PTSD. Nature, doi: 10.1038/s41586-025-09083-y. Epub ahead of print. PMID: 40533550.

Ito R, Lee ACH. (2016). The role of the hippocampus in approach-avoidance conflict decision-making: Evidence from rodent and human studies. Behav Brain Res, 313:345–357

Kaehler ST, Singewald N, Philippu A. (1999). Dependence of serotonin release in the locus coeruleus on dorsal raphe neuronal activity. Naunyn Schmiedebergs Arch Pharmacol, 359(5):386–93

Kaye AP, Krystal JH. (2020). Predictive processing in mental illness: Hierarchical circuitry for perception and trauma. Journal of Abnormal Psychology, 129(6):629–632

Kim EJ, Kong MS, Park SG, Mizumori SJY, Cho J, Kim JJ. (2018). Dynamic coding of predatory information between the prelimbic cortex and lateral amygdala in foraging rats. Sci Adv, 4(4):eaar7328

Kirlic N, Young J, Aupperle RL. (2017). Animal to human translational paradigms relevant for approach avoidance conflict decision making. Behav Res Ther, 96:14–29

Krystal JH, Neumeister A. (2009). Noradrenergic and serotonergic mechanisms in the neurobiology of posttraumatic stress disorder and resilience. Brain Res, 1293:13–23

La-Vu M, Tobias BC, Schuette PJ, Adhikari A (2020). To approach or avoid: An introductory overview of the study of anxiety using rodent assays. Frontiers in Behavioral Neuroscience, 14, 145

Leger L, Descarries L. (1978). Serotonin nerve terminals in the locus coeruleus of adult rat: a radioautographic study. Brain Res, 145(1):1–13

Li L, Feng X, Zhou Z, Zhang H, Shi Q, Lei Z, Shen P, Yang Q, Zhao B, Chen S, Li L, Zhang Y, Wen P, Lu Z, Li X, Xu F, Wang L. (2018). Stress accelerates defensive responses to looming in mice and involves a locus coeruleus-superior colliculus projection. Curr Biol, 28(6):859–871

Li Y, Zhong W, Wang D, Feng Q, Liu Z, Zhou J, Jia C, Hu F, Zeng J, Guo Q, Fu L, Luo M. (2016). Serotonin neurons in the dorsal raphe nucleus encode reward signals. Nat Commun, 7:10503

Liu RJ, Aghajanian GK. (2008). Stress blunts serotonin- and hypocretin-evoked EPSCs in prefrontal cortex: role of corticosterone-mediated apical dendritic atrophy. Proc Natl Acad Sci USA, 105(1):359–64

Liu Z, Zhou J, Li Y, Hu F, Lu Y, Ma M, Feng Q, Zhang JE, Wang D, Zeng J, Bao J, Kim JY, Chen ZF, El Mestikawy S, Luo M. (2014). Dorsal raphe neurons signal reward through 5-HT and glutamate. Neuron, 81(6):1360–1374

Marcinkiewcz CA, Mazzone CM, D’Agostino G, Halladay LR, Hardaway JA, DiBerto JF, Navarro M, Burnham N, Cristiano C, Dorrier CE, Tipton GJ, Ramakrishnan C, Kozicz T, Deisseroth K, Thiele TE, McElligott ZA, Holmes A, Heisler LK, Kash TL. (2016). Serotonin engages an anxiety and fear-promoting circuit in the extended amygdala. Nature, 537(7618):97–101

Marzo A, Totah NK, Neves RM, Logothetis NK, Eschenko O. (2014). Unilateral electrical stimulation of rat locus coeruleus elicits bilateral response of norepinephrine neurons and sustained activation of medial prefrontal cortex. J Neurophysiol, 111(12):2570–88

Massé F, Hascoët M, Dailly E, Bourin M. (2006). Effect of noradrenergic system on the anxiolytic-like effect of DOI (5-HT2A/2C agonists) in the four-plate test. Psychopharmacology (Berl), 183(4):471–81

Mathis A, Mamidanna P, Cury KM, Abe T, Murthy VN, Mathis MW, Bethge M. (2018). DeepLabCut: markerless pose estimation of user-defined body parts with deep learning. Nat Neurosci, 21(9):1281–1289

Miller SM, Marcotulli D, Shen A, Zweifel LS. (2019). Divergent medial amygdala projections regulate approach-avoidance conflict behavior. Nat Neurosci, 22(4):565–575

Nanez J. (1988). Perception of impending collision in 3-to 6-week-old human infants. Infant Behavior and Development, 11(4):447–463

Ortega JE, Mendiguren A, Pineda J, Meana JJ. (2012). Regulation of central noradrenergic activity by 5-HT(3) receptors located in the locus coeruleus of the rat. Neuropharmacology, 62(8):2472–9

Park YG, Sohn CH, Chen R, McCue M, Yun DH, Drummond GT, Ku T, Evans NB, Oak HC, Trieu W, Choi H, Jin X, Lilascharoen V, Wang J, Truttmann MC, Qi HW, Ploegh HL, Golub TR, Chen SC, Frosch MP, Kulik HJ, Lim BK, Chung K. (2018). Protection of tissue physicochemical properties using polyfunctional crosslinkers. Nat Biotechnol, 10.1038/nbt.4281

Pickel VM, Joh TH, Reis DJ. (1977). A serotonergic innervation of noradrenergic neurons in nucleus locus coeruleus: demonstration by immunocytochemical localization of the transmitter specific enzymes tyrosine and tryptophan hydroxylase. Brain Res, 131(2):197–214

Ranjbar-Slamloo Y, Fazlali Z. (2020). Dopamine and Noradrenaline in the Brain; Overlapping or Dissociate Functions? Frontiers in Molecular Neuroscience, 12:334

Ren C, Tao Q. (2020). Neural Circuits Underlying Innate Fear. Adv Exp Med Biol, 1284:1–7

Ren J, Friedmann D, Xiong J, Liu CD, Ferguson BR, Weerakkody T, DeLoach KE, Ran C, Pun A, Sun Y, Weissbourd B, Neve RL, Huguenard J, Horowitz MA, Luo L. (2018). Anatomically Defined and Functionally Distinct Dorsal Raphe Serotonin Sub-systems. Cell, 175(2):472–487.e20

Ritter A, Habusha S, Edut S, Klavir O. (2022). Prefrontal Control of Innate Escape Behavior – A Neural Mechanism of Enhanced Posttraumatic Threat Detection. bioRxiv, 2022.12.21.521361

Salamone JD, Correa M, Yang JH, Rotolo R, Presby R. (2018). Dopamine, effort-based choice, and behavioral economics: Basic and translational research. Frontiers in Behavioral Neuroscience,12:25

Salamone JD, Correa M. (2012) The mysterious motivational functions of mesolimbic dopamine. Neuron, 76(3):470–85

Salay LD, Ishiko N, Huberman AD. (2018). A midline thalamic circuit determines reactions to visual threat. Nature, 557(7704):183–189

Schiff W, Caviness JA, Gibson JJ. (1962). Persistent fear responses in rhesus monkeys to the optical stimulus of “looming”. Science, 136(3520):982–3

Schultz W, Dickinson A. (2000). Neuronal coding of prediction errors. Annu Rev Neurosci, 23:473–500

Schultz W. (2010). Dopamine signals for reward value and risk: basic and recent data. Behav Brain Funct, 6:24

Schultz W. (2016). Dopamine reward prediction error coding. Dialogues Clin Neurosci, 18(1):23–32

Sciolino NR, Hsiang M, Mazzone CM, Wilson LR, Plummer NW, Amin J, Smith KG, McGee CA, Fry SA, Yang CX, Powell JM, Bruchas MR, Kravitz AV, Cushman JD, Krashes MJ, Cui G, Jensen P. (2022). Natural locus coeruleus dynamics during feeding. Sci Adv, 8(33):eabn9134

Shang C, Chen Z, Liu A, Li Y, Zhang J, Qu B, Yan F, Zhang Y, Liu W, Liu Z, Guo X, Li D, Wang Y, Cao P. (2018). Divergent midbrain circuits orchestrate escape and freezing responses to looming stimuli in mice. Nat Commun, 9(1):1232

Shang C, Liu Z, Chen Z, Shi Y, Wang Q, Liu S, Li D, Cao P. (2015). A parvalbumin-positive excitatory visual pathway to trigger fear responses in mice. Science, 348(6242):1472–7

Silva BA, Gross CT, Gräff J. (2016). The neural circuits of innate fear: detection, integration, action, and memorization. Learn Mem, 23(10):544–55

Su Z, Cohen J. (2022). Two types of locus coeruleus norepinephrine neurons drive reinforcement learning. bioRxiv, 2022.12.08.519670

Sun F, Zeng J, Jing M, Zhou J, Feng J, Owen SF, Luo Y, Li F, Wang H, Yamaguchi T, Yong Z, Gao Y, Peng W, Wang L, Zhang S, Du J, Lin D, Xu M, Kreitzer AC, Cui G, Li Y. (2018). A genetically encoded fluorescent sensor enables rapid and specific detection of dopamine in flies, fish, and mice. Cell, 174(2):481–496.e19

Sun F, Zhou J, Dai B, Qian T, Zeng J, Li X, Zhuo Y, Zhang Y, Wang Y, Qian C, Tan K, Feng J, Dong H, Lin D, Cui G, Li Y. (2020). Next-generation GRAB sensors for monitoring dopaminergic activity in vivo. Nat Methods, 17(11):1156–1166

Szabo ST, Blier P. (2001). Serotonin (1A) receptor ligands act on norepinephrine neuron firing through excitatory amino acid and GABA(A) receptors: a microiontophoretic study in the rat locus coeruleus. Synapse, 42(4):203–12

Ungless MA. (2004). Dopamine: the salient issue. Trends Neurosci, 27(12):702–6

van der Weel FR, van der Meer AL. (2009). Seeing it coming: infants’ brain responses to looming danger. Naturwissenschaften, 96(12):1385–91

Vander Weele CM, Siciliano CA, Matthews GA, Namburi P, Izadmehr EM, Espinel IC, Nieh EH, Schut EHS, Padilla-Coreano N, Burgos-Robles A, Chang CJ, Kimchi EY, Beyeler A, Wichmann R, Wildes CP, Tye KM. (2018). Dopamine enhances signal-to-noise ratio in cortical-brainstem encoding of aversive stimuli. Nature, 563(7731):397–401

Varazzani C, San-Galli A, Gilardeau S, Bouret S. (2015). Noradrenaline and dopamine neurons in the reward/effort trade-off: a direct electrophysiological comparison in behaving monkeys. J Neurosci, 35(20):7866–77

Ventura R, Latagliata EC, Morrone C, La Mela I, Puglisi-Allegra S. (2008). Prefrontal norepinephrine determines attribution of “high” motivational salience. PLoS One, 3(8):e3044

Wan J, Peng W, Li X, Qian T, Song K, Zeng J, Deng F, Hao S, Feng J, Zhang P, Zhang Y, Zou J, Pan S, Shin M, Venton BJ, Zhu JJ, Jing M, Xu M, Li Y. (2021). A genetically encoded sensor for measuring serotonin dynamics. Nat Neurosci, 24(5):746–752

Wei P, Liu N, Zhang Z, Liu X, Tang Y, He X, Wu B, Zhou Z, Liu Y, Li J, Zhang Y, Zhou X, Xu L, Chen L, Bi G, Hu X, Xu F, Wang L. (2015). Processing of visually evoked innate fear by a non-canonical thalamic pathway. Nat Commun, 6:6756

Yilmaz M, Meister M. (2013). Rapid innate defensive responses of mice to looming visual stimuli. Curr Biol, 23(20):2011–5

Yu AJ, Dayan P. (2005). Uncertainty, neuromodulation, and attention. Neuron, 46(4):681–92

Zald DH, Treadway MT. (2017). Reward processing, neuroeconomics, and psychopathology. Annu Rev Clin Psychol, 13:471–495

Zambetti PR, Schuessler BP, Kim JJ. (2019). Sex Differences in Foraging Rats to Naturalistic Aerial Predator Stimuli. iScience, 16:442–452

Zambetti PR, Schuessler BP, Lecamp BE, Shin A, Kim EJ, Kim JJ. (2022). Ecological analysis of Pavlovian fear conditioning in rats. Commun Biol, 5(1):830

Zhao X, Liu M, Cang J. (2014). Visual cortex modulates the magnitude but not the selectivity of looming-evoked responses in the superior colliculus of awake mice. Neuron, 84(1):202–213

Zhou Z, Liu X, Chen S, Zhang Z, Liu Y, Montardy Q, Tang Y, Wei P, Liu N, Li L, Song R, Lai J, He X, Chen C, Bi G, Feng G, Xu F, Wang L. (2019). A VTA GABAergic neural circuit mediates visually evoked innate defensive responses. Neuron, 103(3):473–488

Zhu Y, Dewell RB, Wang H, Gabbiani F. (2018). Pre-synaptic muscarinic excitation enhances the discrimination of looming stimuli in a collision-detection neuron. Cell Rep, 23(8):2365–2378

Zhuo Y, Luo B, Yi X, Dong H, Wan J, Cai R, Williams JT, Qian T, Campbell MG, Miao X, Li B, Wei Y, Li G, Wang H, Zheng Y, Watabe-Uchida M, Li Y. Improved dual-color GRAB sensors for monitoring dopaminergic activity in vivo. *bioRxiv*, 2023.08.24.554559; doi: 10.1101/2023.08.24.554559

